# Linking Genetic Risk to Disease-Relevant Cellular States via Metacell-Informed Modeling with ICePop

**DOI:** 10.64898/2026.04.01.715877

**Authors:** Hao Yuan, Aishwarya Mandava, Kewalin Samart, Julia Ganz, Arjun Krishnan

## Abstract

Genome-wide association studies (GWAS) have implicated thousands of loci in complex diseases, but translating these population-level signals into specific cellular contexts remains a central challenge. Integrating GWAS with single-cell transcriptomics data has enabled systematic identification of disease-relevant cell types, yet existing methods face a fundamental tradeoff: approaches like seismic that optimized for statistical power operate at the annotated cell-type level and miss heterogeneous disease signals concentrated in specific cellular states, while single-cell-resolution approaches like scDRS that capture such heterogeneity often lack sufficient power to detect subtle associations. Here we present ICePop (**I**nformative **Ce**ll **Pop**ulations), a framework that resolves this tradeoff by performing disease-cell type association at metacell resolution, thus achieving statistical power comparable to cell-type-level methods while detecting heterogeneous disease signals within cell types. In simulations against seismic and scDRS, ICePop maintains appropriate false positive rates and demonstrates superior power when disease effects are concentrated in cellular subpopulations. Applied to Tabula Muris across 81 traits and 120 cell types, ICePop identifies 1,684 disease-cell type associations, including the preferential vulnerability of differentiated gut epithelial cells in ulcerative colitis and loss of cell identity in immune-stressed lung capillary endothelial cells underlying their association with lung function. Clustering diseases by metacell association profiles reveals groupings that diverge from genetic risk-based clustering, including separation of blood cell count traits from immune diseases despite shared genetic architecture, reflecting differences in cellular rather than genetic etiology. In autism spectrum disorder, ICePop identifies preferential enrichment of genetic risk in specific enteric neuron subtypes, implicating dysfunction of the enteric nervous system in gastrointestinal comorbidities. ICePop’s resolution of disease-relevant cell states within annotated cell types enables generation of testable, cell-state-specific hypotheses about disease mechanisms and therapeutic targets.

## Introduction

Genome-wide association studies (GWAS) have identified thousands of disease-associated loci, yet translating these population-level signals into specific cellular contexts remains a major challenge. A central question is which cell types and cellular states mediate genetic risk [1]. Resolving this question is critical for understanding disease mechanisms and prioritizing therapeutic targets.

To address this problem, many computational frameworks integrate GWAS signals with single-cell gene expression data to infer disease-associated cell types. A dominant strategy aggregates SNP-level associations into gene-level disease scores, typically using MAGMA [2], and then tests whether genes that are highly associated with disease are enriched among cell-type-specific expression programs. A representative method in this category is scDRS [3] defining putative disease-associated genes by selecting top-ranked genes based on MAGMA z-scores. For each individual cell, it samples control genes matching expression distribution of disease genes and gene set size, then tests whether the putative disease genes have higher expression than the matched controls. Statistical testing is performed at the single-cell level and subsequently aggregated to the cell-type level. This single-cell resolution enables scDRS to capture within-cell-type heterogeneity. scDRS implicitly assumes that disease-associated genes should exhibit higher overall expression than control genes, which can be overly restrictive. In practice, many disease-associated genes are not highly expressed but instead have high specificity for particular cellular contexts. By emphasizing absolute expression rather than specificity, such approaches may miss subtle, context-dependent signals, especially for complex traits where genetic effects are modest and distributed [4].

A more recent representative method is seismic [5], which improves statistical power by quantifying disease-cell type associations through linear regression between gene-level disease score (MAGMA z-scores) and cell-type expression specificity. Because a positive association reflects concordance between genetic risk and expression specificity—rather than requiring uniformly high expression of top-ranked disease genes—seismic is more sensitive to specificity-driven disease signals. Accordingly, compared to scDRS, seismic demonstrated increased sensitivity in causal simulations while maintaining good calibration in null settings [5]. However, this gain in power comes with an important limitation: seismic performs association testing only at the cell-type level and does not provide a mechanism to resolve how disease risk varies across heterogeneous cellular subpopulations within a cell type.

Cellular heterogeneity in disease association is pervasive and represents a critical component of disease mechanisms and therapeutic target prioritization [6, 7]. A canonical example is the association between microglia and Alzheimer’s disease. Only a specific subpopulation—disease-associated microglia—is enriched around amyloid plaques, a hallmark of Alzheimer’s pathology, where they contribute to synaptic disruption and neurodegeneration. In contrast, the majority of homeostatic microglia do not contribute directly to disease progression [8, 9, 10]. This example illustrates that disease risk is often concentrated in specific cellular states rather than uniformly distributed across an entire cell type, underscoring the need for methods that can resolve disease-relevant heterogeneity without sacrificing statistical power.

A key challenge in resolving disease-cell type associations while accounting for cellular heterogeneity is the tradeoff between resolution and statistical stability. Analyses at the level of individual cells offer fine-grained resolution but are severely affected by technical noise and dropout, whereas analyses at the level of annotated cell types improve robustness at the cost of obscuring heterogeneous disease signals within a cell type. Metacells provide an intermediate representation by aggregating transcriptionally similar cells into homogeneous groups, reducing technical noise while preserving biologically meaningful heterogeneity [11]. This representation offers a natural unit for modeling disease associations that vary across cellular states within a cell type.

Building on this idea, we developed **I**nformed **Ce**ll **Pop**ulation (ICePop), a framework that models disease-cell associations by leveraging metacells as the fundamental unit to capture cellular heterogeneity. ICePop extends specificity score-based regression to the metacell level, enabling association testing at an intermediate resolution that balances noise reduction with preservation of biological variation. To address dependencies arising from similar expression profiles across metacells—which can otherwise inflate cell type-level associations—we use a covariance-aware permutation null distribution that accounts for the correlation structure among metacells, and thereby controls inflation of cell-type-level associations. ICePop further applies a weighted FDR framework to prioritize strongly associated metacells within each cell type, facilitating detection of heterogeneous disease signals. In addition, it incorporates a DFBETAS-based approach to identify influential genes by quantifying metacell-level contributions and emphasizing those from associated metacells.

To enhance interpretability, ICePop generates an interactive report for exploring disease-cell type heterogeneity and the underlying genes. Together, these innovations provide a robust and interpretable framework for resolving disease associations across complex cellular landscapes.

Here, we evaluate the performance of ICePop in detecting heterogeneous disease signals and demonstrate its utility and robustness through several analyses. First, we designed a novel simulation framework that preserves realistic gene expression distributions and cellular diversity, and performed systematic evaluation under both null and causal simulation settings. Second, we applied ICePop to identify disease-associated cell types across 81 traits and diseases using the Tabula Muris (TM) FACS dataset [12], thereby assessing its performance in large-scale disease association screening. Third, we compared metacell association profiles across traits and diseases to identify shared and distinct cellular mechanisms. Finally, we characterized heterogeneity in disease associations across gut cell populations implicated in autism spectrum disorder (ASD).

## Results

### Overview of ICePop for disease-cell type association and heterogeneity

As shown in **Fig. 1A**, ICePop began by aggregating single cells into metacells, which represent transcriptionally homogeneous cellular states. We generated metacells using MetaQ [13], a scalable framework designed for metacell generation in large single-cell datasets. For each gene and each metacell, we computed an expression specificity score that quantifies how selectively a gene is expressed in a metacell relative to other metacells. In parallel, gene-level disease association statistics were derived from GWAS summary statistics using MAGMA [2]. These gene-level association scores were then regressed against metacell-specific expression specificity scores to obtain metacell-level disease association coefficients.

**Figure 1:**
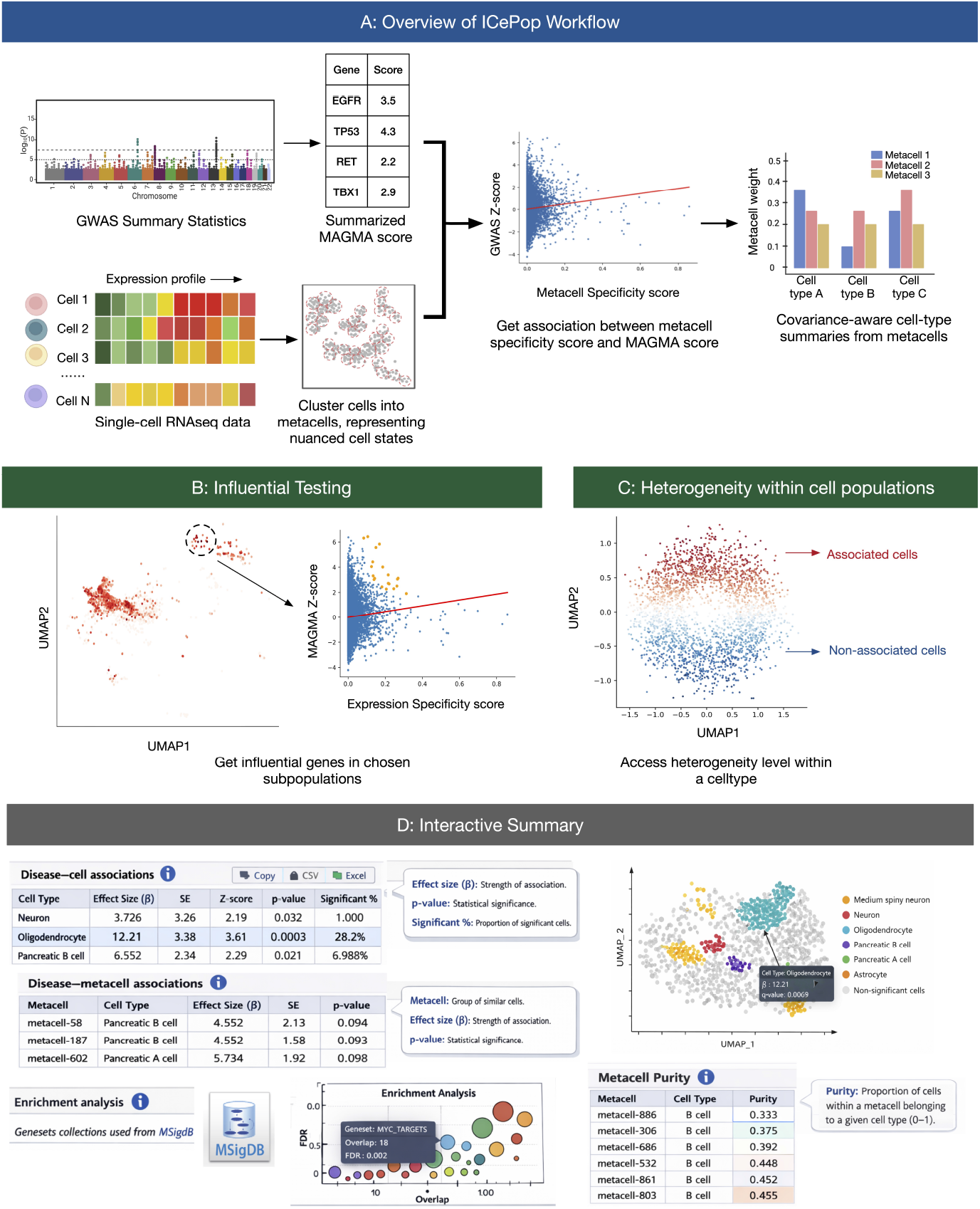
(A) ICePop Workflow: ICePop integrates GWAS summary statistics and single-cell transcriptomic data to identify disease-cell type associations. GWAS signals are summarized from SNPs to gene-level association scores using MAGMA, while single-cell expression profiles are aggregated into metacells that capture cell states. For each metacell, an expression specificity score is computed and regressed against the corresponding disease association score to quantify disease-metacell similarity. Metacell-level association statistics are then aggregated to the cell-type level using a covariance-aware framework that accounts for expression similarity among metacells, yielding cell-type-level association scores and significance. **(B) Influential testing:** For downstream analysis, influential testing identifies genes driving associations within user-selected cell populations (circled), with influential genes highlighted in orange. **(C) Heterogeneity within cell populations:** Heterogeneity level of association within each cell type is assessed. **(D) Interactive Summary:** At the end of the pipeline, a fully interactive HTML report is generated to provide a comprehensive summary of all downstream analyses including disease association and metacell statistics, and hypergeometric test performed using curated geneset collections from MSigDB, highlighting biologically relevant pathways and transcriptional programs enriched in disease-associated cell types. The report includes interactive tables, sortable columns, searchable fields, downloadable outputs (CSV/Excel), and dynamic visualizations (e.g., UMAP embeddings and enrichment bubble plots), enabling intuitive exploration and interpretation of results.

To infer cell-type-level disease associations, coefficients of all metacells within each cell type were aggregated using a weighted averaging scheme that emphasizes metacells with high cell-type purity (the proportion of cells within a metacell belonging to a given cell type) and strong disease association. Because transcriptionally adjacent metacells tend to exhibit correlated expression profiles and regression coefficients, we used a covariance-aware null distribution that preserves the correlation structure among metacells when estimating cell-type-level associations. To further control false positives, we restricted the contribution of low-quality metacells to disease-cell type associations based on their size and purity (see Methods).

We implemented influence diagnostics based on DFBETAS [14] to prioritize genes that disproportionately contribute to disease-cell type associations. These diagnostics explicitly incorporated the same weighting scheme used in the metacell-to-cell type aggregation step, thereby emphasizing gene contributions arising from metacells that are strongly associated with the disease and represent relevant cellular subpopulations within each cell type. In addition, ICePop supports the computation of disease associations for user-defined cell groups, enabling flexible exploration beyond predefined cell-type annotations.

To assess cellular heterogeneity within cell types, we applied weighted Benjamini-Hochberg procedures [15] to identify significantly associated metacells while accounting for contribution of each metacell to its corresponding cell type. Significance assignments at the metacell level were then propagated to individual cells, allowing us to quantify the proportion of significantly associated cells within each cell type and thereby characterize within-cell-type heterogeneity.

For each disease, ICePop generates an interactive report that allows users to explore associated cell populations, contributing genes, and enriched biological processes. Together, these features provide a comprehensive and interpretable view of disease-cell type relationships, capturing within-cell-type heterogeneity and offering insight into the contextual roles of genes in specific disease-cell type combinations.

### ICePop is well calibrated and powerful under heterogeneous disease-cell type associations

We compared the performance of ICePop to scDRS [3], a widely used method for identifying disease-cell-type associations, and also seismic [5], a recent method shown to outperform earlier approaches such as FUMA [16] and S-MAGMA [17]. We conducted comprehensive and systematic null simulation experiments to assess whether all methods are well calibrated, that is, whether they appropriately control type I error under the null hypothesis. These null simulations were performed using randomly subsampled cells from TM FACS and permuted gene-disease association scores (see Methods).

We first assessed the calibration of disease-metacell associations. Metacells of all sizes (75, 50, and 30 cells per metacell representing large, medium, and small metacells) exhibited comparable and well-controlled type I error rates, indicating that metacell size does not affect calibration (**Supplementary Fig. 1 a-c**). At the cell-type level, we assessed calibration by computing calibration performance per cell type across all runs. Type I error was generally well controlled for all methods and three metacell sizes when compared across cell types (**Fig. 2A**). ICePop showed slight inflation at extreme quantiles, a pattern that remained consistent when aggregating associations across different metacell sizes (**Supplementary Fig. 2a-c**). A similar slight inflation was also observed for seismic. Further stratification by cell-type size revealed no systematic differences in calibration across cell-type size bins for any metacell size in ICePop (**Supplementary Fig. 3a-c**) or across methods (**Supplementary Fig. 4a-b**).

**Figure 2:**
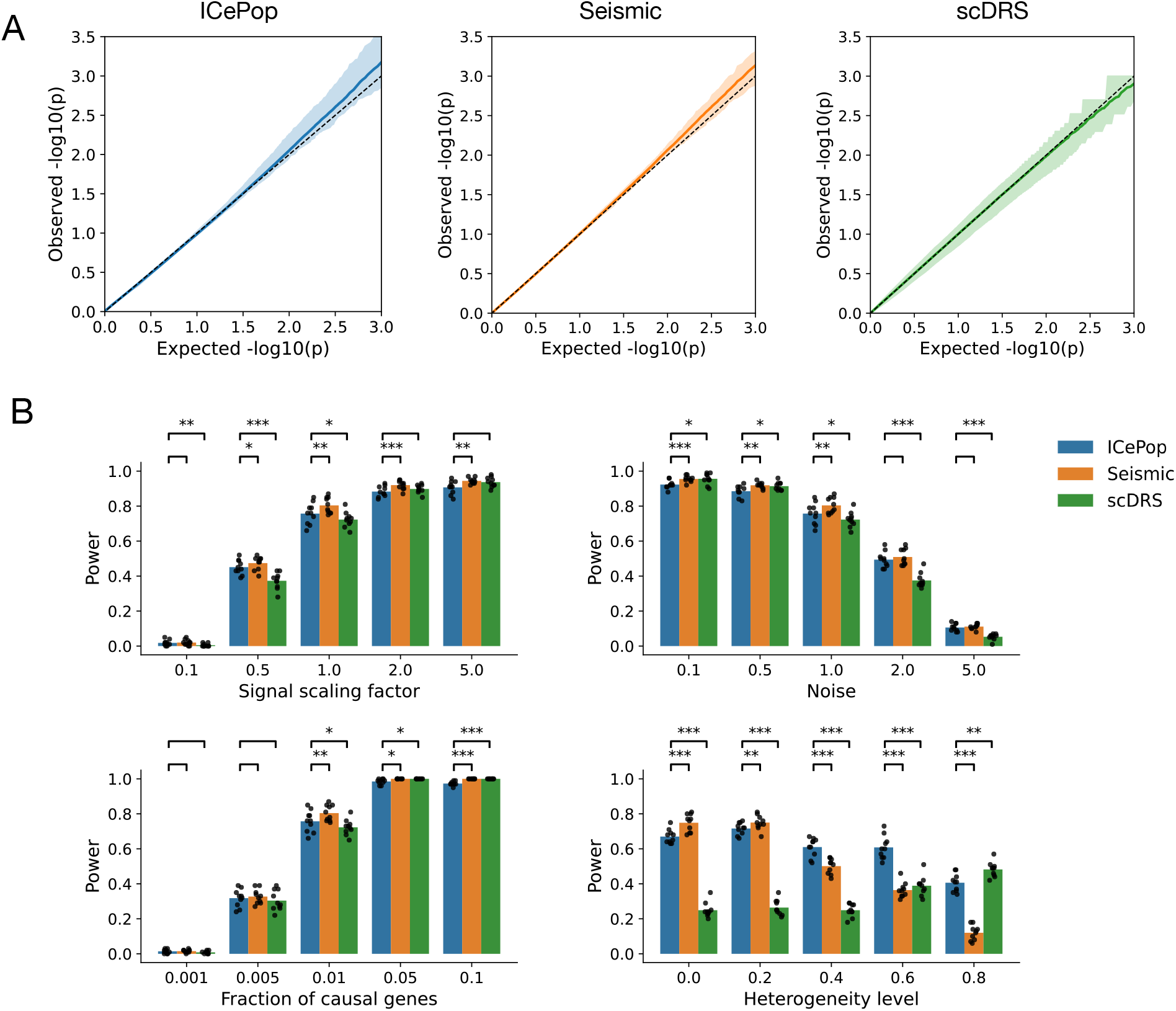
(A) Quantile-quantile (QQ) plots for null simulations. Null distributions were generated by permuting MAGMA z-scores for 10,000 runs. The x-axis shows expected p-values under the null, and the y-axis shows observed p-values for disease-cell type associations computed between permuted MAGMA z-scores and subsets of 10,000 cells sampled from TM FACS data. Shaded regions indicate variability across cell types, bounded by the 5th and 95th percentiles. The dashed line denotes the expected null distribution, and the solid line represents the average observed − log_10_(*p*-value) across cell types. **(B) Power comparisons for disease-cell type association under four simulation parameters**. MAGMA z-scores were synthetically generated based on cell-type-specific gene expression, with signal strength controlled by fraction of causal genes, noise level, signal scaling factor, and heterogeneity level (see Methods). Analyses were conducted on a subset of 10,000 cells from the Tabula Muris FACS dataset. Each bar represents the proportion of simulations (out of 100 independent runs) in which the true causal cell type was correctly identified as significant (FDR ≤ 0.1). Black dots indicate the variations of power estimates across 10 repetitions. Significance levels of difference between ICePop and seismic or scDRS are denoted as * (*p* < 0.05), ** (*p* < 0.01), and *** (*p* < 0.001). In each panel, one parameter is varied while the remaining three are held fixed. Unless otherwise specified, default parameter values are fraction of causal genes = 0.01, noise = 1.0, signal scaling factor = 1.0, and heterogeneity level = 0.0.

We next evaluated statistical power under scenarios in which true disease-cell type associations were present. Our simulation framework assumes that a cell type is disease-associated when disease-risk genes overlap with genes specifically expressed in that cell type. Based on this principle, we generated association signals using cell-type-specific expression patterns and designated a subset of specifically expressed genes as causal, with non-causal genes modeled as noise. We varied key parameters, including the fraction of causal genes, noise level, and signal scaling factor, to represent realistic disease signals. To preserve the empirical distribution of GWAS statistics, simulated gene effects were mapped onto gene-disease scores within randomly selected traits. For each parameter setting, we performed 100 simulations and defined power as the proportion of runs in which the target cell type reached FDR of 0.1. The entire procedure was repeated 10 times per setting to assess variability (see Methods).

To simulate heterogeneous disease associations within a cell type, we introduced a heterogeneity parameter controlling the proportion of truly disease-associated cells. Specifically, we constructed a synthetic cell type by selecting a localized patch of transcriptionally similar cells as the disease-associated subpopulation and a separate, transcriptionally distant patch as the non-associated subpopulation. The heterogeneity level was defined as the proportion of non-associated cells within the synthetic cell type, such that higher heterogeneity corresponds to a smaller fraction of truly disease-associated cells. We then quantified differences in gene expression specificity between associated and non-associated subpopulations to derive disease scores, thereby generating disease signals enriched within the associated subpopulation (see Methods).

First, we assessed the effect of power on ICePop when aggregating signal from varying metacell sizes. Across simulations varying key parameters, metacell size had minimal impact on overall power (**Supplementary Fig. 5a-c**). Under high heterogeneity level, we observed a slight decrease in power for larger metacells, but performance remained broadly comparable across sizes (**Supplementary Fig. 5d**). Stratifying results by cell-type size yielded a similar pattern: metacell sizes of 30 and 50 showed higher power at higher heterogeneity level (**Supplementary Fig. 6d**), whereas no systematic differences were observed under other parameter settings (**Supplementary Fig. 6a-c**). Although smaller metacells showed slight advantages in high heterogeneity scenarios, larger metacells are expected to provide more robust expression specificity estimates due to increased signal aggregation. Given its comparable performance across simulation settings, we selected a metacell size of 75 for subsequent analyses, beginning with the comparison of ICePop to seismic and scDRS.

Across simulations varying the fraction of causal genes, noise level, and signal strength, ICePop and seismic showed similar performance. scDRS performed well under strong signals but had reduced power as signals weakened, such as with fewer causal genes, higher noise, or lower scaling factors (**Fig. 2B**). When heterogeneity in disease signal increased, ICePop consistently outperformed seismic, with the performance gap widening at higher heterogeneity levels. In contrast, scDRS showed reduced power at low heterogeneity but improved as heterogeneity increased, approaching ICePop’s performance. This pattern is consistent with the design of scDRS to detect subpopulation-specific signals (**Fig. 2B**).

Stratifying performance by cell-type size revealed nuanced differences across methods. Overall, ICePop remained comparable to seismic, although seismic showed a slight advantage in very small cell types (≤ 61 cells), particularly under unrealistically strong signal settings (signal scaling factor = 5.0, noise = 0.1, and causal gene fraction = 0.1). This likely reflects the use of a metacell size of 75 in ICePop; when a cell type contains fewer cells than the average target metacell size, metacells that containing some cells from neighbor cell types could slightly reduce power. However, performance remained highly comparable, and this difference disappeared in larger cell types (*≥* 124 cells) (**Supplementary Fig. 7a-c**). Because these simulations were performed on a subset of TM FACS dataset, real datasets are expected to contain larger cell-type populations, where ICePop achieved performance comparable to seismic. Across size bins, increasing heterogeneity substantially reduced the power of seismic, whereas ICePop maintained more stable performance across heterogeneity levels, especially in larger cell types (≥ 127 cells) (**Supplementary Fig. 7d)**.

Consistent with the overall trend of performance before cell-type size stratification, scDRS showed reduced power under weaker signal settings across cell-type size bins, including lower scaling factors, higher noise, or smaller causal gene fractions. This reduction was more pronounced in larger cell types (≥ 163 cells) (**Supplementary Fig. 7a-c**). Performance of scDRS improved as heterogeneity increased. However, from low (0.2) to moderately high heterogeneity levels (0.6), ICePop was consistently comparable to or better than scDRS. At extreme heterogeneity, 0.8, ICePop performed better in larger cell types (≥ 160 cells), whereas scDRS had an advantage in smaller cell types (≤ 152 cells). The lower power of ICePop in small cell types under extreme heterogeneity may be due to a single metacell in the cell type that cannot capture heterogeneity. However, biologically, strong heterogeneity is more likely to occur in larger cell populations, where ICePop showed a clear power advantage (**Supplementary Fig. 7d**).

### ICePop detects meaningful disease-cell type association from real data, particularly with cell-type heterogeneity

To evaluate the performance of ICePop in identifying disease-cell type associations across a broad range of cell types and diseases in real data, we applied the method to the TM FACS dataset. We curated traits spanning seven major disease categories and included 120 cell types in the analysis, together encompassing 21 tissues and 81 traits (**Supplementary Data 1**). For ICePop, we used a metacell size of 75 cells to construct disease associations, as justified in **Supplementary Note 3**. ICePop identified 1,684 significant disease-cell type associations in total, a number lower yet similar in magnitude to seismic (n=1,786), whereas scDRS detected substantially fewer (n=903). Associations identified by ICePop shared the most with those detected by seismic (n=690) (**Fig. 3B**).

**Figure 3:**
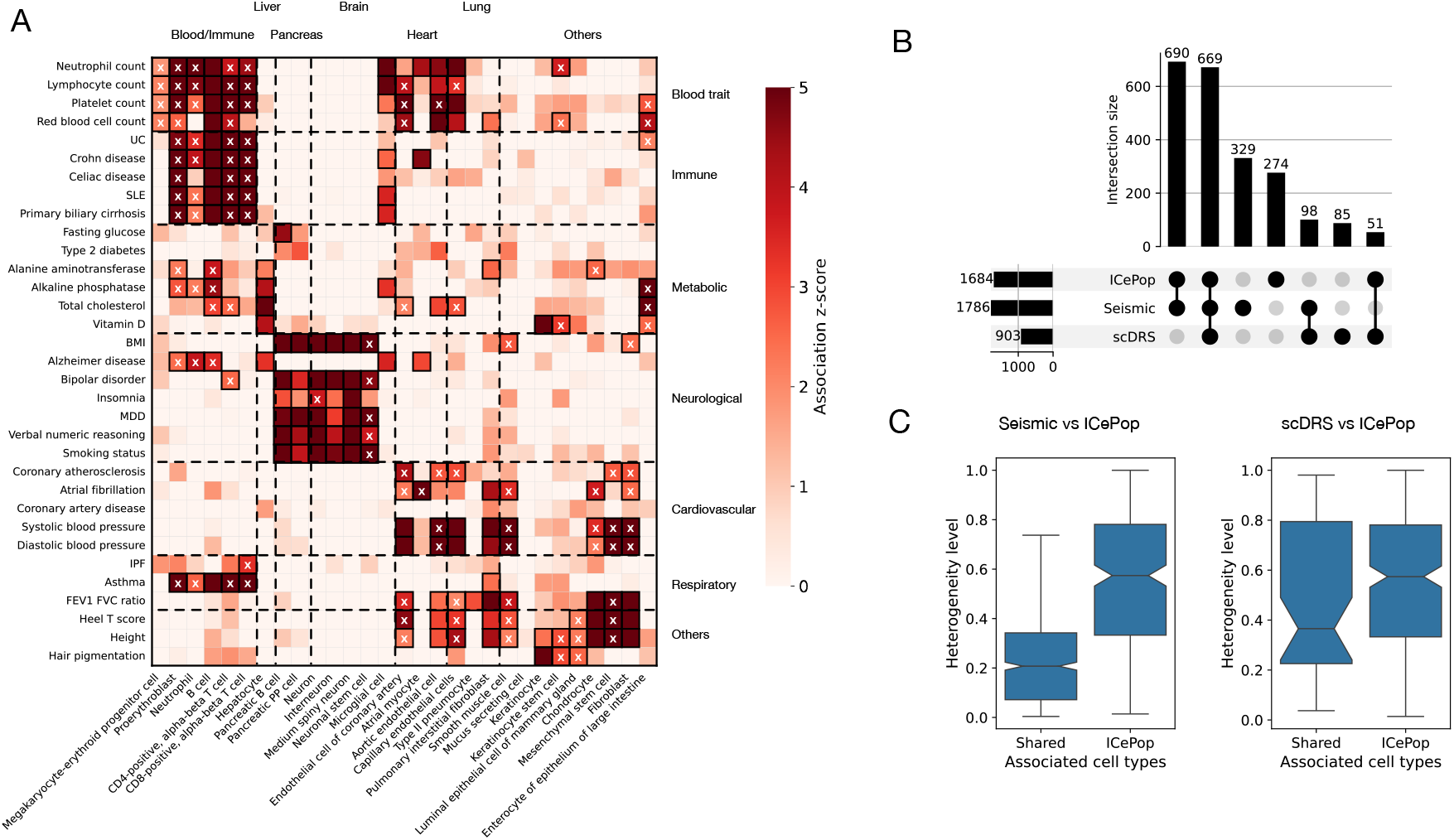
(A) Heatmap of ICePop disease-cell-type association results for 33 representative traits and 29 cell types from the Tabula Muris FACS dataset. Rows correspond to diseases or traits, and columns correspond to cell types, grouped into seven disease categories and seven tissues or organs. Colors indicate cell-type association z-scores, with negative values truncated to zero. Black rectangles denote significant associations, and cross symbols indicate heterogeneity of disease association, defined as significant association (FDR ≤ 0.1) present in more than 20% but fewer than 80% of cells within a cell type. Trait abbreviations are ulcerative colitis (UC), systemic lupus erythematosus (SLE), body mass index (BMI), major depressive disorder (MDD), and idiopathic pulmonary fibrosis (IPF). Association results for all 81 traits and 120 cell types are provided in Supplementary Data 2. **(B) UpSet plot summarizing shared and unique disease-cell type associations identified from TM FACS data**. Left bar plots indicate the total number of associations per method, while top intersection bars show number of shared and method-specific associations. **(C) Characterization of heterogeneity among shared and unique associations identified by ICePop and other methods**. The left panel compares cell types jointly identified by Seismic and ICePop (left boxplot) with those uniquely identified by ICePop (right boxplot). The right panel presents the analogous comparison between scDRS and ICePop. Only cell types containing at least two significant metacells are included. The y-axis represents the level of within-cell-type heterogeneity of disease association, defined as 1 - the proportion of significant metacells within that cell type; higher values indicate greater heterogeneity.

Associations identified by all three methods (ICePop, seismic, and scDRS; n=669) recapitulated well-established disease-cell type relationships. These included blood cells with hematological traits, immune diseases with immune cell types, neurological disorders with neuronal cells, metabolic traits with hepatocytes, and height with chondrocytes (**Fig. 3A, Supplementary Fig. 8 and 9**). The concordance across methods indicates strong and robust disease-cell type signals that are consistently detectable regardless of an-alytical framework. Associations identified by both ICePop and seismic (n=690) included many non-obvious links that were less frequently detected by scDRS (**Fig. 3A, Supplementary Fig. 8**). These include links between pancreatic cells and neurological traits, supported by the fact that insulin-secreting pancreatic cells relay systemic metabolic information (e.g., glucose levels) to the brain through sensory neurons, supporting biologically meaningful cross-organ associations [18]. ICePop and seismic consistently co-identified an association between hair pigmentation and keratinocyte stem cells, with FDR=0.026 for ICePop and 0.001 for seismic (**Fig. 3A, Supplementary Fig. 8**). Keratinocytes, as the primary structural cells of the hair shaft, receive and retain melanin produced by melanocytes, thereby involving determination of pigmentation [19].

A smaller subset of associations was uniquely shared between ICePop and scDRS (n=51) and tended to reflect heterogeneous associations. For instance, both ICePop and scDRS found that blood pressure-related traits, such as diastolic blood pressure, was heterogeneously associated with aortic endothelial cells (ICePop: 20.56% significant cells, scDRS: heterogeneity level FDR=0.053) (**Fig. 3A, Supplementary Fig. 9**). Plaque formation on endothelial cells, including aortic endothelial cells, is closely linked to changes in diastolic blood pressure [20]. However, plaque does not affect all endothelial cells uniformly. Instead, it preferentially develops in regions of disturbed blood flow, such as vascular branch points, rather than in streamlined vascular segments [21]. Similarly, ICePop and scDRS found that the FEV1/FVC ratio (the ratio of forced expiratory volume in one second to forced vital capacity, a measure of lung function) was associated with lung capillary cells (ICePop: 42.18% significant cells, scDRS: heterogeneity level FDR=0.014) (**Fig. 3A, Supplementary Fig. 9**). The heterogeneity underlying this association is discussed in greater detail below.

We further characterized ICePop-specific associations (n=274), typically corresponding to cases where just a fraction of cells within a cell type are associated with a disease, many of which reflecting biologically meaningful heterogeneity. For example, ICePop identified a heterogeneous association between atrial fibrillation and fibroblasts (FDR=0.055, 38.50% significant cells), consistent with evidence that only activated fibroblasts differentiate into myofibroblasts involved in fibrillation pathogenesis [22] (**Fig. 3A**). ICePop also identified an association between ulcerative colitis and large-intestinal enterocytes (FDR=0.071, 28.05% significant cells), consistent with heterogeneous enterocyte responses during intestinal inflammation (**Fig. 3A**) (discussed below).

Associations shared between ICePop and seismic usually occurred in cell types where most cells were associated with the trait. In contrast, ICePop-specific associations had a much lower fraction of associated cells, indicating that ICePop can detect heterogeneity within cell types (**Fig. 3C**). Associations shared between ICePop and scDRS also showed low fractions of associated cells, suggesting that both methods can detect heterogeneous signals. However, ICePop-specific associations generally showed higher overall heterogeneity compared to scDRS (**Fig. 3C**). When assessing shared associations stratified by tissue, we observed a similar overall trend: seismic showed greater overlap with ICePop than scDRS did. Tissues such as brain, immune, blood, and liver exhibited a high proportion of shared disease-cell type associations across methods, indicating robust and relatively homogeneous signals. In contrast, tissues including skin and large intestine showed lower overlap and a higher fraction of ICePop-specific associations, suggesting that disease-cell type relationships in these tissues are more heterogeneous and therefore preferentially captured by ICePop (**Supplementary Fig. 10a-b**).

A substantial number of cases were uniquely identified by seismic. Stratifying associations by cell type size revealed that 191 of 383 such cases occurred in the smallest cell type size bin (4-72 cells) (**Supplementary Fig. 11a**), a regime in which cell types are smaller than the average metacell size (75). However, this range also includes very small cell types, which may yield unstable and less reliable estimates. In contrast, ICePop tended to detect more associations in larger cell type size bin (≥ 1,429 cells) (**Supplementary Fig. 11e**), suggesting that heterogeneous disease-associated cell states within larger populations are better captured by ICePop. Additional examples of seismic-specific identifications are discussed in **Supplementary Note 4**.

### ICePop uncovers nuanced disease-associated cell state changes within cell types

We next investigated cases exhibiting substantial heterogeneity within cell types in TM FACS. We focused on scenarios involving subtler disease-associated cell state changes, allowing us to evaluate whether ICePop can detect nuanced within-cell-type variation.

One illustrative example is the association between large intestine enterocytes and ulcerative colitis (UC).

Enterocytes line the colonic mucosa and serve as the primary epithelial barrier and absorptive surface [23]. In UC, dysregulated immune responses can damage gut epithelia including enterocytes, leading to immunestress-mediated damage such as epithelial erosion, ulceration, and increased intestinal permeability [24, 25, 26]. However, epithelial damage is not spatially uniform, suggesting that not all enterocytes are equally vulnerable to immune stress [27].

Consistent with these facts, we observed that the association between enterocytes and UC was heterogeneous rather than being uniform across the cell type (FDR=0.071, %28.05 significant cells). To further characterize this pattern, we assigned individual cells to disease-associated or non-associated groups (see Methods). Associated and non-associated cells largely overlapped in UMAP space, indicating that the transcriptional differences were subtle and did not reflect discrete cell-type separation (**Fig. 4A**). This was further supported by broad expression of canonical enterocyte markers (*CDH1* [28], *CA1* [29], *KRT20* [30]) across both groups (**Supplementary Fig. 12b**), suggesting that the observed differences represent nuanced cell state variation within the cell type.

**Figure 4:**
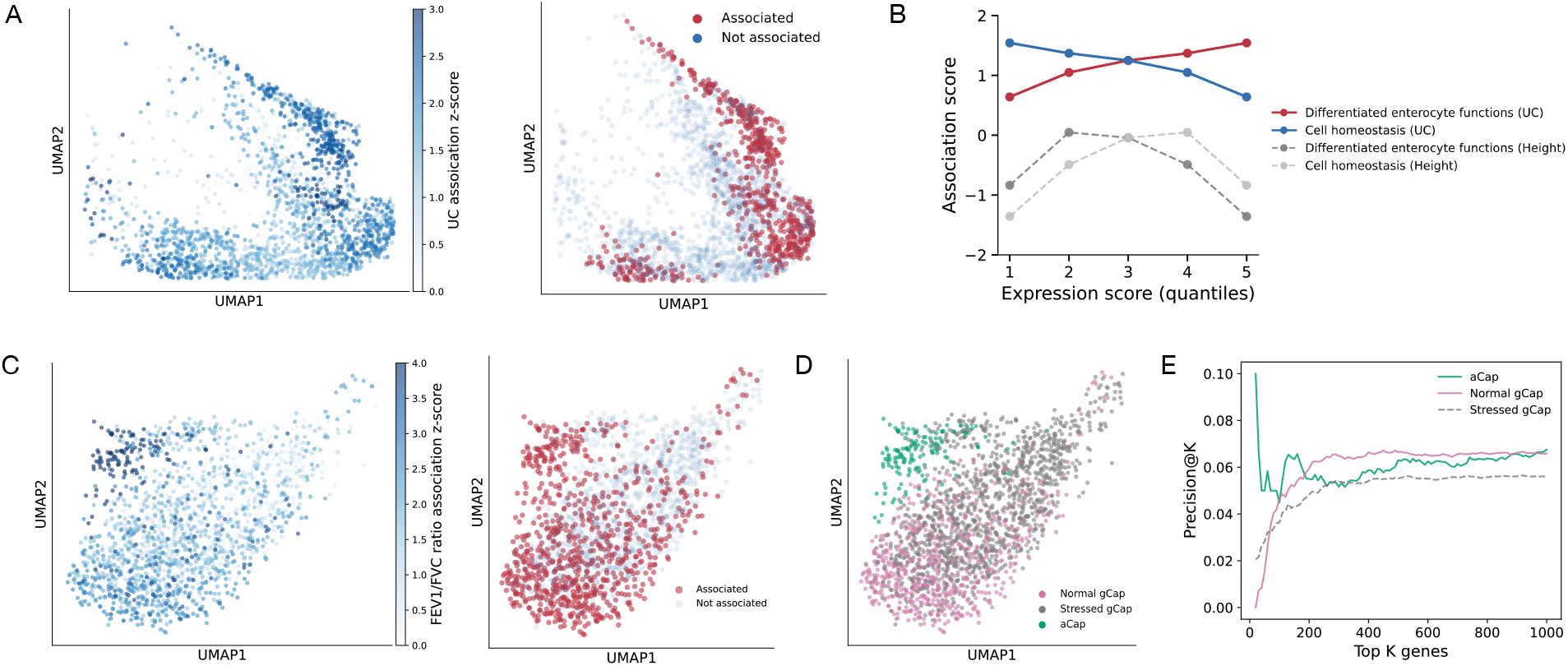
(A) Heterogeneity of UC association within large-intestinal enterocytes. Left panel shows metacell-level association scores projected onto individual enterocytes. The displayed cells represent 96.93% of the total population (full view shown in Supplementary Fig. 12a). Right panel shows cells classified as significantly associated (red) or not associated (blue) with UC based on metacell-level FDR ≤ 0.1. **(B) Relationship between gene expression programs and UC association**. The x-axis shows quantiles of relative expression strength for two gene programs: differentiated enterocyte functions (red) and cell homeostasis (blue) (see Methods). Cells are grouped into five quantile bins based on the relative expression of program-specific expression. The y-axis shows the corresponding mean metacell-level association score for UC across cells. Height was included as a negative control, where the y-axis represents the metacell-level z-scores for height association using the same expression-based binning. **(C) Heterogeneity of FEV1/FVC association within lung capillary endothelial cells**. Left panel shows metacell-level association scores projected onto individual cells (94.85% of cells shown; full view in Supplementary Fig. 14a). Right panel shows cells classified as significantly associated (red) or not associated (blue) with FEV1/FVC. **(D) Loss of cell identity in lung capillary endothelial cells under immune stress**. Left panel shows annotation of cell states based on marker expression (see Supplementary Fig. 14b and 15). Right panel shows enrichment of FEV1/FVC related genes in specifically expressed genes from each cell-state group. The x-axis represents the top k specifically expressed genes from each group. The y-axis shows precision at k, defined as the proportion of genes within the top 5% of MAGMA z-scores for FEV1/FVC, averaged across metacells within each group.

Previous studies have shown that mature enterocytes are more vulnerable to apoptosis and immune stress than progenitor cells, particularly under endoplasmic reticulum (ER) stress conditions [31, 32]. Based on this, we asked whether UC-associated enterocytes are more differentiated, whereas non-associated cells may be less differentiated and exhibit stronger cell homeostasis under stress. To answer this question, for each cell, we calculated the relative expression of genes annotated to sets of Gene Ontology Biological Processes, one representing differentiated enterocyte functions (e.g., ion transport and water absorption) and the other representing cellular homeostasis programs, including ER stress responses, apoptosis, and epithelial differentiation (see Methods). We found that UC-associated enterocytes exhibited higher relative expression of processes linked to differentiated epithelial functions. In contrast, non-associated cells showed higher expression of processes related to cellular homeostasis. Height, which has minimal association to enterocytes, did not show bias toward either processes (**Fig. 4B**). We further examined well-known markers of differentiated enterocytes. Ion transport genes such as *SLC26A3* [33] and *ATP12A* [34], as well as the water channel gene *AQP8* [35], were more highly expressed in UC-associated cells. In contrast, non-associated cells expressed higher levels of genes related to cellular homeostasis and metabolic maintenance, including *PRKG2* [36], *ATP5IF1* [37], and *RNF186* [38] (**Supplementary Fig. 12b and 13a-i**). Together, these results suggest that UC-associated enterocytes are enriched for mature epithelial functions, whereas non-associated cells maintain gene expression programs linked to cell differentiation and homeostasis.

Another illustrative example is the association between lung capillary endothelial cells and FEV1/FVC ratio, a clinical measure of airway obstruction and lung function. Capillary endothelial dysfunction has been implicated in reduced lung function and progressive fibrosis [39]. In our analysis, we observed substantial within-cell type heterogeneity in capillary endothelial cells, with 42.18% of cells at significant levels. On UMAP, capillary endothelial cells are separated into two major populations corresponding to aerocytes (aCap) and general capillary endothelial cells (gCap), according to marker expression (**Supplementary Fig. 14b**). While aCap cells were broadly disease-associated, the gCap population exhibited clear heterogeneity, with a subset showing weaker association signals (**Fig. 4C**).

Prior mouse studies have shown that endothelial cells in aged lungs with sustained fibrosis exhibit reduced expression of endothelial identity genes and increased inflammatory programs [39]. We therefore hypothesized that loss of cell identity may reduce detectable disease-cell type association signals, as association frameworks rely on overlap between disease-risk genes and cell population-specific expression signatures. Consistent with this hypothesis, the less-associated gCap state retained general gCap features but displayed elevated immune-stress signatures, including increased expression of interferon genes (*IFITM3*) [40] and immune regulatory genes (*TMSB4X*) [41] (**Supplementary Fig. 14b and 15g-h**), indicating that this gCap subset exists in a stress-like state, even though TM data were derived from healthy, wild-type mice. This stressed gCap population also exhibited reduced expression of canonical gCap markers (*TEK, APLNR*) [42] (**Supplementary Fig. 14b and 15i-k**). We next identified specifically expressed genes for each meta-cell using the metacell specificity score (see Methods) and tested whether genes strongly associated with FEV1/FVC ratio were enriched among these specifically expressed genes. We found that aCap and normal gCap metacells showed higher enrichment compared to stressed gCap metacells (**Fig. 4D and E**). This pattern supports the idea that the observed heterogeneity is related to the erosion of gCap identity in the stressed state.

### ICePop revealed shared cellular mechanism among diseases

Understanding shared mechanisms across diseases is essential for elucidating disease pathophysiology, improving diagnosis, and guiding treatment design. While GWAS provides important insights, similarities based solely on shared genetic risk may be limited due to substantial genetic heterogeneity across complex traits [43]. Estimating similarities based on shared cellular programs instead offers a more nuanced, mechanistic, and biologically interpretable view of relationships between diseases. We systematically characterized disease similarity using metacell-level disease association profiles and compared these patterns with gene-based risk similarity. We further clustered the disease-disease similarity network to identify key differences between cellular association patterns and genetic risk similarity (see Methods).

We observed substantial overlap between metacell-based and genetic risk-based clustering. For example, red blood cell traits (e.g., mean corpuscular hemoglobin, red blood cell distribution width, and red blood cell count) clustered together in both frameworks. Neurological traits and diseases also formed consistent groups. Lung function traits (FEV1/FVC ratio and lung FVC) clustered together, as did skeletal traits (heel T score and height) (**Fig. 5A, Supplementary Fig. 16**). These concordant groupings suggest that these traits not only share genetic risk architecture but also share similar cellular mechanisms.

**Figure 5:**
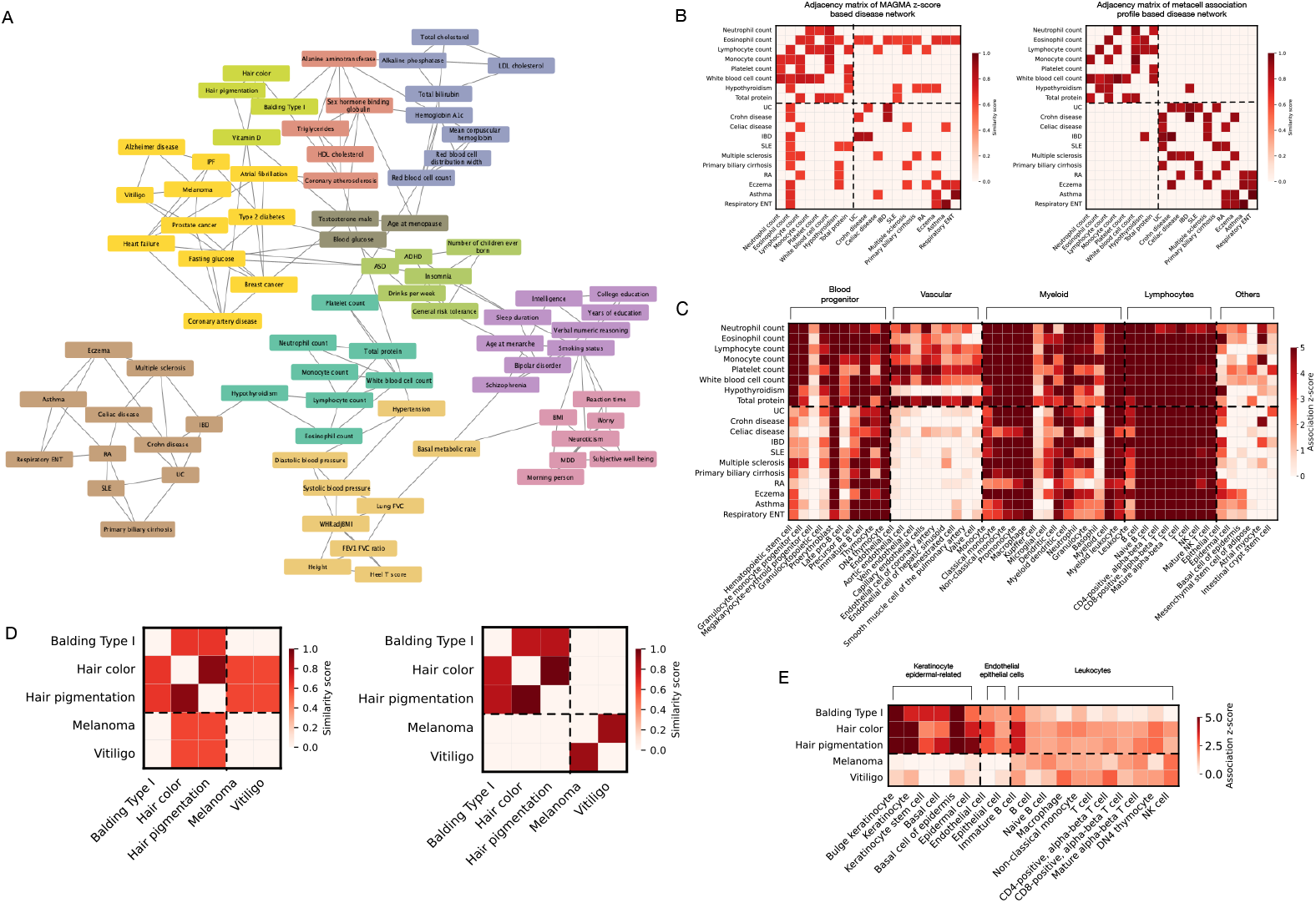
(A) Disease-disease similarity graph based on metacell profile similarity, highlighting shared cellular mechanisms. The graph is constructed from similarities among metacell-level z-score profiles using a k-nearest-neighbor graph, where k=3. Louvain clustering is applied at a resolution of 2.5, with each color representing a distinct cluster of traits or diseases. **(B) Subnetwork of the disease-disease network for immune diseases and leukocyte count-related traits**. Both panels display subnetworks represented as adjacency matrices. Left panel shows a network constructed using MAGMA z-scores across traits. Rows and columns represent diseases, and color intensity indicates edge weight (pairwise similarity). Right plot shows corresponding subnetwork for the same set of traits constructed using similarity of metacell-level z-score profiles. **(C) Cell type-trait associations for immune diseases and leukocyte count-related traits**. Each row represents a trait and each column represents a cell type. Color indicates ICePop cell type association scores. Score below zero is truncated. **(D) Subnetwork of the disease-disease network for pigment-related traits**. Network is constructed and visualized as in (B), using either MAGMA z-scores (left) or metacell profile similarity (right). **(E) Cell type-trait associations for pigment-related traits**. As in (C), rows represent disease traits and columns represent cell types. Color indicates zero-truncated ICePop cell type association scores.

However, differences also emerged. One major divergence involved myeloid and lymphocyte count traits. In genetic risk-based clustering, these traits are closely connected with immune diseases such as ulcerative colitis (UC), systemic lupus erythematosus (SLE), and multiple sclerosis due to shared immune-related genetic signals (**Fig. 5B, Supplementary Fig. 16**). In contrast, metacell-based clustering separated myeloid and lymphocyte cell count traits from immune diseases (**Fig. 5B**). Myeloid and lymphocyte counts were more strongly associated with upstream progenitor populations, including hematopoietic stem cells (shared origin), granulocytopoietic cells (myeloid branch), and late pro-B cells (lymphoid branch), as well as vascular endothelial cells. These progenitor and vascular populations were not strongly associated with immune diseases (**Fig. 5C**). This suggests that although immune diseases and blood cell counts share genetic architecture, their cellular manifestations differ.

Another illustrative example is pigmentation-related traits. ICePop grouped hair traits (balding type I, hair color, and hair pigmentation) together and separated from melanoma and vitiligo (**Fig. 5D**), reflecting strong involvement of keratinocytes [44], epidermal cells [45] and endothelial cells [46] in hair growth (**Fig. 5E**). In the genetic risk-based comparison, melanoma and vitiligo showed some similarity to hair pigmentation traits (**Fig. 5D, Supplementary Fig. 16**). This suggests overlap in pigmentation-related risk genes, even though these conditions involve different cell types in how they manifest. Overall, our method provides insight into the cellular mechanisms underlying disease similarity, adding an additional layer of biologically interpretable disease comparisons beyond shared genetic risk.

### ICePop revealed heterogeneous enteric neuron association to Autism spectrum disorder

Autism spectrum disorder (ASD) is primarily classified as a neurodevelopmental condition, but many individuals also experience gastrointestinal (GI) symptoms, including abdominal pain, diarrhea, and constipation [47, 48]. These comorbidities implicate gut cell dysfunction. Recent studies have reported associations between ASD genetic risk and gut cell populations, including enteric neurons [49] and intestinal epithelial cells [50, 51]. However, GI symptom vary widely across individuals with ASD, suggesting that the affected cell populations and the downstream programs perturbed within them may be context-dependent and heterogeneous [51]. To further investigate heterogenous association of enteric cells to ASD, we applied ICePop to a colonic cell dataset to prioritize cell populations that are more associated with ASD and to characterize heterogeneity within these cell types.

We analyzed mouse colon single-cell data from Drokhlyansky et al. [49], encompassing 22 annotated cell types and capturing the broad cellular diversity in the mouse colon (**Supplementary Fig. 17a**). We used GWAS summary statistics of ASD from Pedersen et al. [52]. ICePop identified enteric neurons as the most strongly ASD-associated population (**Supplementary Fig. 17b and c**). Within the enteric neurons, sensory neurons, secretomotor/vasodilator neurons, and inhibitory motor neurons showed the strongest association (**Fig. 6A, Supplementary Table 1, Supplementary Data 3**). These findings are consistent with prior reports linking these neuronal subtypes to GI dysfunction in ASD [53, 54, 55]. In contrast, excitatory motor neurons showed borderline association (p = 0.062, FDR = 0.274). For non-neuronal cell type, enteroendocrine cells, which have also been suggested to play a role in ASD-related gut dysfunction [56], similarly exhibited marginal signals (p = 0.061, FDR = 0.274). Detection from seismic and scDRS showed drastic differences.

**Figure 6:**
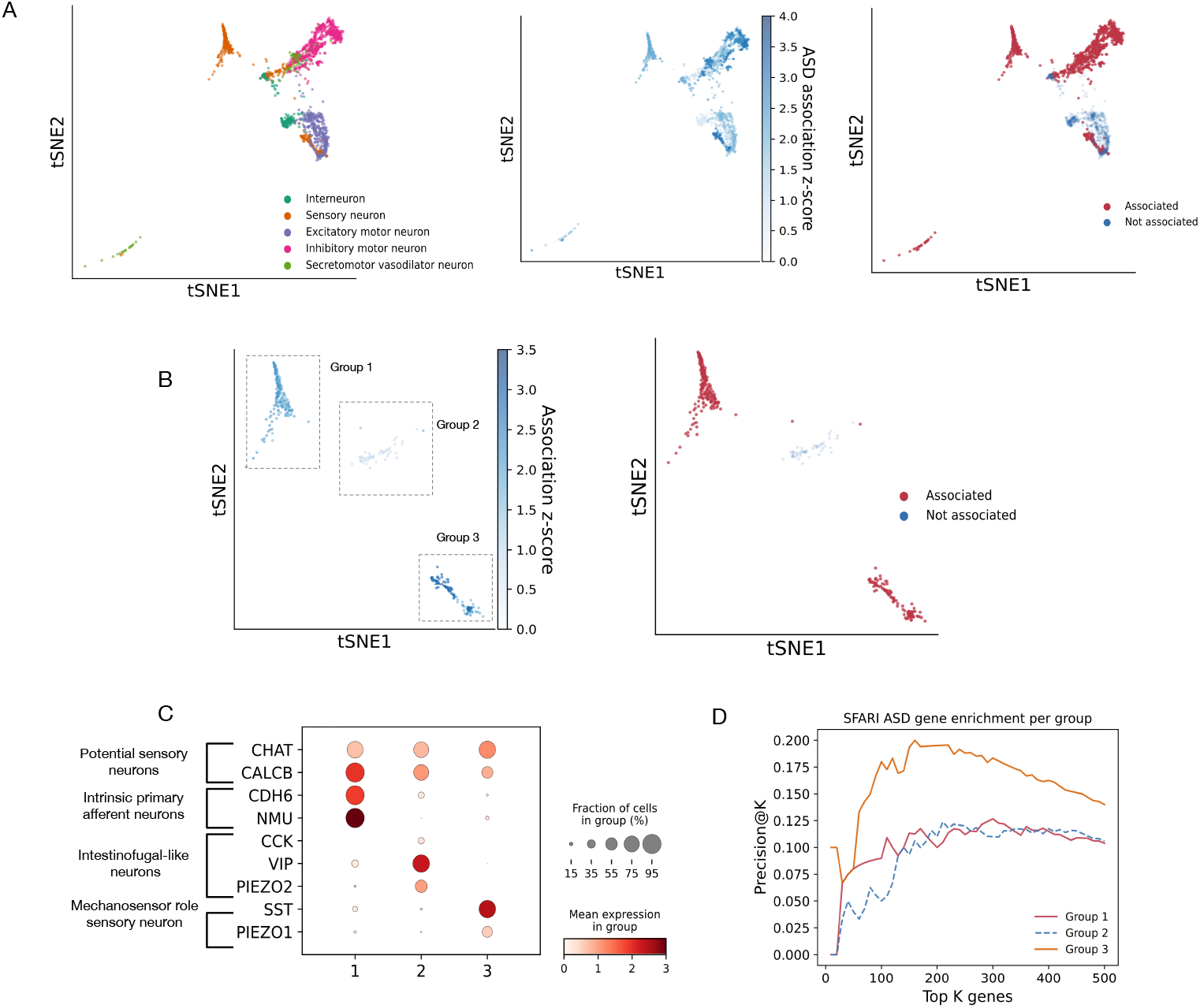
(A) Heterogeneity of ASD association across enteric neuron cell types. The left panel shows the distribution of enteric neuron cell types, the middle panel shows metacell-level ASD association scores projected onto individual cells, and the right panel shows significantly associated enteric neurons identified by propagating metacell-level significance to individual cells using metacell-level FDR (≤ 0.1). **(B) Heterogeneity of ASD association within enteric sensory neurons**. The left panel shows metacell-level ASD association scores projected onto individual cells in a tSNE embedding (cells shown correspond to 98.95% of enteric sensory neurons, full tSNE shown in Supplementary Fig. 18a). Cell groups are highlighted in the plot. Right panels shows significant associated cells identified by propagating metacell-level significance to individual cells using metacell-level FDR (≤ 0.1). **(C) Expression of marker genes indicating distinct subgroups of enteric sensory neurons**. The x-axis shows the cell groups defined in panel B. The y-axis lists marker genes representing distinct subtypes of enteric sensory neurons. Dot color indicates log transformed expression level, and dot size reflects the fraction of cells expressing each gene within a group. **(D) Enrichment of high-confidence SFARI ASD genes among top influential genes in cell groups**. The plot shows the enrichment of high-confidence SFARI genes (SFARI gene score ≤ 2.0) in top influential genes that contribute to disease-cell group associations. The x-axis indicates the top K influential genes ranked within each subgroup, and the y-axis shows precision at K. Each line corresponds to a distinct enteric sensory neuron subgroup.

Seismic detected only secretomotor/vasodilator neurons, whereas scDRS identified no significant associations (**Supplementary Table 1, Supplementary Data 3**).

We observed substantial heterogeneity among ASD-associated enteric neurons. Within each cell type, only a subset showed association with ASD, including 80.0% of sensory neurons, 55.24% of secretomotor/vasodilator neurons, and 63.76% of inhibitory motor neurons. (**Supplementary Data 3**). We focused on dissecting heterogeneity within sensory neurons to determine whether specific subpopulations exhibited elevated vulnerability to ASD-associated genetic risk. Sensory neurons fell into three major groups, corresponding to six metacells (**Fig. 6A, Supplementary Fig. 18b**). Canonical markers indicated that all three groups represent sensory neuronal populations, as defined by expression of *CHAT* and *CALCB* [57, 58]. Group 1 displayed co-expression of *CDH6* and *NMU*, consistent with intrinsic primary afferent neurons (IPANs) [57]. Expression of weak yet specific *CCK* further suggests a role in transmitting signals from the gut wall to pre-vertebral ganglia in group 2 [59]. Group 3 showed elevated *PIEZO1* expression, indicating mechanosensory function [60] (**Fig. 6C**). Group 2 exhibited weaker association signals, suggesting a comparatively reduced role in ASD (**Fig. 6B**).

We hypothesized that Groups 1 and 3 are preferentially associated with ASD because a greater number of genuine ASD risk genes are specifically expressed in these subpopulations, thereby contributing more strongly to their disease-cell group association signals. To test this, we extracted high-confidence ASD genes from the Simons Foundation Autism Research Initiative (SFARI) database [61] (SFARI gene score ≤ 2.0). We then computed aggregated gene influence scores within each neuronal group to quantify the contribution of individual genes to disease association, applying the same framework used to identify influential genes at the cell-type level to the neuronal group level (see Methods). Groups 1 and 3 showed stronger enrichment for high-confidence ASD genes compared to Group 2, supporting their increased vulnerability to ASD (**Fig. 6D**). Additionally, we examined genes known to disrupt sensory neuron development or function, including those affecting neural crest migration (*CDH8, CDH2*) [62], synaptic function (*CNTNAP2*) [63], and serotonin sensing (*HTR3A*) [64]. These genes were more broadly expressed in Groups 1 and 3 than in Group 2, further reinforcing their preferential association with ASD (**Supplementary Fig. 18c**). Altogether, ICePop provides insight into the heterogeneous nature of ASD associations within enteric neurons and reveals substantial heterogeneity even within a single cell type.

## Discussion

Thus far, disease-cell type association and within-cell-type heterogeneity have largely been treated as separate problems requiring distinct analytical frameworks. Methods such as scDRS [3] capture heterogeneity but often sacrifice statistical power, whereas approaches like seismic [5] maximize power at the cost of collapsing within-cell-type variation. In this study, we introduce ICePop, which resolves this trade-off by selecting an appropriate intermediate unit of analysis—metacells—rather than attempting to address both challenges at either the single-cell or cell-type level.

Metacells have been shown to effectively mitigate the fundamental tension between resolution and statistical stability in single-cell analysis. While single-cell measurements are prone to dropout and noise, and cell-type-level aggregation can obscure meaningful variation, metacells provide an intermediate representation that amplifies biological signal while preserving local structure [11]. This representation has proven compatible with a wide range of single-cell analyses, including clustering, differential expression, imputation, and RNA velocity [65, 66, 13]. Building on this foundation, we extend the metacell paradigm to disease-cell association analysis. By operating at a granularity that better matches the biological scale of disease processes, ICePop uncovers fine-grained heterogeneity within cell types, capturing continuous cellular state transitions that shape disease association patterns.

As one of the first applications of ICePop, we used it to uncover the cellular basis of ulcerative colitis (UC). Our analysis shows that UC genetic risk is not uniformly distributed across epithelial cells, but is instead enriched in terminally differentiated states. While prior studies have established the central role of epithelial cells in UC pathogenesis [67, 68, 69, 70], they largely focused on broad cell types. Our results further refined this view by showing that genetic risk is strongly dependent on cellular state within the epithelial lineage, with enrichment in cells performing terminal functions such as ion transport and absorption, rather than in proliferative or stress-resistant states. This suggests that epithelial vulnerability in UC is concentrated in the differentiated layer and has implications for experimental modeling: organoid systems enriched for progenitor-like states may underestimate the genetic contribution to UC epithelial pathology.

Another application analyzing the association of FEV1/FVC ratio (measure of airway obstruction and lung function) with lung capillary cells highlights a distinct aspect of cellular heterogeneity. Here, even in healthy tissue, a subset of cells exhibit an immune-stressed state characterized by reduced expression of canonical cell identity markers. This finding suggests that potential disease-relevant cellular states may exist prior to overt pathology. By explicitly modeling such heterogeneity, ICePop can detect these pre-existing, potentially disease-associated states, offering opportunities for early biomarker discovery and prevention-oriented research.

Our analysis of mouse colonic cells in the context of autism spectrum disorder (ASD) reveals that, although the association of enteric neurons with ASD is previously known [54], the risk is not uniformly distributed. We observe the strongest associations in sensory neurons. Within sensory neurons, mechanosensory *PIEZO1* - expressing neurons and intrinsic primary afferent neurons (IPANs) show stronger enrichment, consistent with known enteric nervous system (ENS) biology: IPANs initiate gut activity via chemical and mechanical cues [59, 71], and their disruption impairs sensing and downstream motility [63], while loss of *PIEZO1* is linked to slowed peristalsis [60]. Regulatory circuits are also implicated, with enrichment in secretomotor/vasodilator neurons and inhibitory motor neurons. Secretomotor circuits regulate epithelial fluid secretion, while inhibitory motor neurons control muscle relaxation [72]; impairment in both may contribute to constipation in ASD. These results indicate that ASD risk primarily disrupts upstream sensory pathways, with additional effects on regulatory circuits for gut motility. Together, these findings provide a concrete mechanistic hypothesis for how ASD affects specific enteric neuron groups, facilitating further investigation into the mechanisms underlying ASD-associated gut dysfunction.

A broad application of ICePop to a wide range of traits and diseases shows that, by integrating cellular context, clustering traits/diseases based on their perturbed metacells reveals distinct patterns compared to those solely based on genetic risk. For example, blood cell count traits cluster with immune diseases based on genetic risk due to shared immune genetics, but separate out based on metacell-based clustering because these traits reflect progenitor and vascular programs, whereas immune diseases are driven by mature effector states—consistent with hematopoietic differentiation trajectories [73]. Similarly, the clustering of pigmentation traits illustrates that shared genetic architecture does not necessarily imply shared cellular etiology: hair pigmentation involves keratinocyte and epithelial programs involving hair pigment growth and retention, whereas melanoma and vitiligo do not. By capturing these distinctions, ICePop prioritizes downstream effector cell types over shared upstream genetic signals, revealing biologically meaningful differences that are not apparent from GWAS alone, thus improving disease stratification and mechanistic interpretation.

The assumption that diseases manifest in a cell-type-specific manner underlies specificity-based methods such as ICePop and seismic. When gene risk of diseases is broadly distributed across genes widely expressed in the tissue and therefore having limited cell-type specificity, the association signal may be difficult to detect. In such cases, specificity-based methods may systematically underperform regardless of resolution, as no meaningful specificity can be captured. Under these conditions, reduced power should be interpreted as an absence of detectable cell-type-specific enrichment within the current dataset, rather than a failure of the method. In these scenarios, using datasets with greater cellular diversity or adopting alternative analytical strategies may be more appropriate.

Metacell construction is a critical component of ICePop, as they define the resolution at which disease-cell associations are evaluated. While users can adopt different metacell construction methods, these approaches vary substantially in their properties, including the internal homogeneity of metacells, sensitivity to rare cell types, and computational efficiency. So, the choice of algorithm should be guided by the specific goals and constraints of the analysis. For example, Metacell-2 [74] applies stricter criteria during construction and can exclude low-quality or unassignable cells, resulting in more compact and homogeneous metacells. However, this stringency may come at the cost of discarding cells from rare populations. In contrast, methods such as SEACells [66] and MetaQ [13] retain all cells, making them more appropriate when rare cell types may contribute to disease associations, albeit potentially at the expense of producing looser metacells that may mix signals from different cell types. Computational scalability is another key consideration. Methods that rely on k-nearest neighbor (KNN) graphs can become prohibitively expensive for large datasets. In this context, MetaQ is currently one of the few approaches capable of scaling to very large single-cell datasets (e.g., >200,000 cells), whereas other methods may become computationally infeasible. Therefore, in this study, we use MetaQ as the default method due to its strong balance between metacell quality and scalability for large datasets, with the latter enabling broad survey-style application of ICePop to many diseases and large single-cell atlases.

The absence of a robust gold standard for disease-cell type associations is the field’s most constraining technical gap, impacting all methods including ICePop. Developing a rigorous and less biased benchmarking framework is necessary to fill this gap. Such a framework should go beyond easily detectable, well-known associations (e.g., neurons with neurological diseases or immune cells with immune disorders) and instead capture more subtle and heterogeneous disease-cell relationships. One potential strategy is to leverage case-control single-cell datasets to identify cell populations showing differential expression or abundance, and use these as data-driven proxies for disease-associated cell types. Constructing such a resource should be a community priority, which would benefit not only ICePop but the entire class of GWAS-to-single-cell integration methods.

Short of high-quality detailed gold-standards, this sub-field relies on simulations for model specification and evaluation. However, current simulation approaches for disease-cell type association do not provide a shared framework that generates data suitable for all classes of GWAS-to-single-cell integration methods. This includes not only methods based on summarized gene-level risk, but also SNP-level heritability enrichment approaches such as CELLECT [75] and sc-linker [76]. Developing a unified simulation framework that supports both gene- and SNP-level models would enable more consistent and fair comparisons across methods.

Finally, current approaches identify disease-associated cell types and prioritize contributing genes but lack mechanistic interpretation of underlying regulatory programs. A key step forward is to map disease-associated signals within relevant cell types to mechanism-aware frameworks such as PhenoPLIER [77], which can reveal coordinated gene programs and regulatory modules. In parallel, network-based approaches offer complementary insight by modeling interactions between disease and associated cell population, either through mapping disease-cell association contributing genes onto functional interaction networks (e.g., STRING [78]) or building co-expression [79] or regulatory networks [80] from disease-associated cell populations.

Overall, by building around metacells as the unit for detecting disease-associated cell populations, ICePop enables the investigation of pervasive heterogeneity in disease associations within cell types, opening the way for a more nuanced understanding of how diseases manifest in specific cellular contexts, and opportunities for early biomarker discovery and prevention-oriented research.

## Method

### ICePop framework

#### Clustering cells to metacells

We used MetaQ [13] to cluster cells into metacells. MetaQ employed an autoencoder framework, but instead of compressing expression profiles into continuous embeddings, it assigned each cell to a finite set of embeddings with each embedding representing a metacell. MetaQ was run using default parameters, except for the target number of metacells, which was defined as the total number of cells divided by the expected number of cells per metacell. We initially set the expected metacell size to 75 cells, a value empirically validated by Persad et al. [66] as providing a reasonable balance between resolution and robustness. To assess the impact of metacell size on performance, we further reduced the expected metacell size to 50 and 30 cells.

We selected MetaQ over the widely used SEACells [66] due to scalability concerns. SEACells constructs a K-nearest neighbor (KNN) graph of cells and applies archetypal matrix factorization to define metacells - both of which are computationally intensive, especially for large datasets. In our benchmarking, SEACells took approximately 12 hours to process a dataset of 110,000 cells on a single Intel Xeon CPU. In contrast, MetaQ completed the same task in just 30 minutes on a V100 GPU, demonstrating far greater efficiency and scalability. This performance makes MetaQ a more practical choice for large-scale single-cell data analysis.

### Calculating metacell specificity score

We adapted the cell-type specificity score from seismic [5] to calculate a metacell-specificity score. The score integrates two components: expression specificity 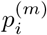 and expression coverage 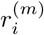. Expression specificity is defined as:

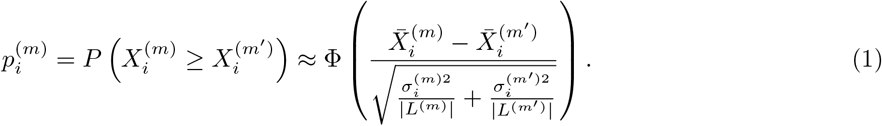

Here, 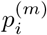 represents the probability that the expression level of gene *i* in metacell *m*, denoted 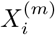, is greater than or equal to its expression in all other metacells *m*′, denoted 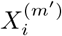, which is further can be interpreted as comparing average of expression in 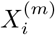 and 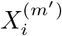, while normalized by standard deviation of gene *i* in metacell group 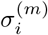 and rest of the cell 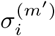 and also size of the metacell group *L*^(*m*)^ and *L*^(*m*′)^. Φ(*·*) is the standard normal CDF.

Expression coverage is defined as:

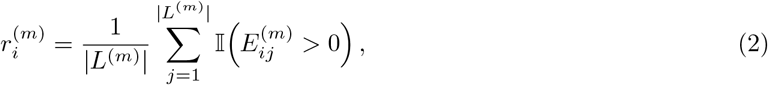

which is essentially percentage of metacell *m* that have positive expression of gene *i*. Finally, the metacell specificity score is calculated as:

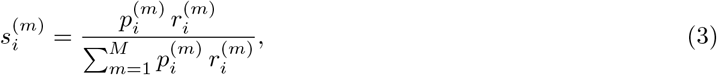

where *M* is the number of metacells. Specificity score is calculated for all genes *i* and metacells *m* in the dataset.

#### Calculating metacell-disease association

Follow idea from seismic [5], we define the metacell-level association for metacell *m* using an ordinary least squares (OLS) linear regression between gene-level GWAS summary scores and metacell specificity scores, and only genes that have MAGMA z-score and also exist in the scRNA-seq dataset are considered.

Let **z** ∈ ℝ^*G*^ denote the vector of MAGMA z-scores across *G* genes, and let 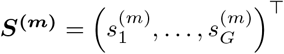 denote the corresponding vector of metacell *m* specificity scores. The regression model is given by:

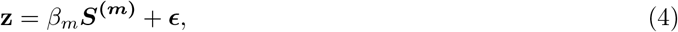

where *β*_*m*_ quantifies the strength of association between metacell *m* and the disease, and ***ϵ*** denotes the residual error term. Both **z** and ***S***^**(*m*)**^ are centered prior to regression, and therefore no intercept term is included. The estimated coefficient 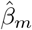 and its corresponding standard error 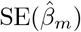 are obtained from the OLS fit.

### Aggregating metacell-level association to cell type level

Cell-type-level association effects are then obtained by computing a weighted average of the selected metacell-level effects, where weights are a combination of cell type purity and statistical significance of metacell to disease association.

For a given metacell *m* and cell type *c*, cell-type purity *π*_*c,m*_ is defined as the proportion of cells in metacell *m* that belong to cell type *c*. The statistical significance of metacell to disease association *α*_*m*_ is incorporated via a one-sided normal cumulative distribution function:

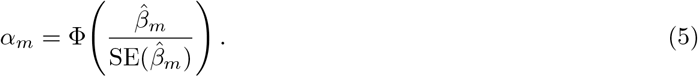

Metacells with very low cell-type purity or very small size are inherently low quality and can introduce spurious associations. To mitigate this, we set the cell-type purity *π*_*c,m*_ of a metacell *m* to zero if its size *n*_*m*_ < 25. This threshold is motivated by modeling the number of cells from cell type *c* within a metacell as a binomial process. Let *k*_*c,m*_ denote the number of cells of type *c* in metacell *m*, with total size *n*_*m*_. We place a Beta prior on purity, *π*_*c,m*_ ~ Beta(*a, b*), and use a non-informative prior *a* = *b* = 1. The posterior distribution is then

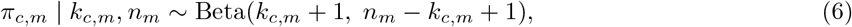

with posterior variance

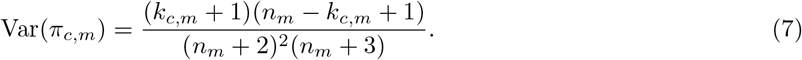

As *n*_*m*_ increases, the posterior variance decreases and begins to stabilize around *n*_*m*_ *≈* 25, irrespective of cell type purity, supporting our minimum size threshold (**Supplementary Fig. 19**).

In addition, we regularize *π*_*c,m*_ to zero for metacells with *π*_*c,m*_ < 0.2. Interpreting 1 − *π*_*c,m*_ as the probability that a metacell does not belong to cell type *c*, this corresponds to a liberal threshold to exclude highly ambiguous metacells.

For every cell type *c*, regularized weights are normalized to sum of one across metacells:

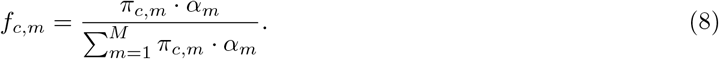

Let 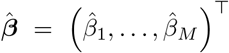 denotes the vector of estimated metacell-level association effects. Let ***f***_*c*_ = (*f*_*c*,1_, …, *f*_*c,M*_)^*⊤*^ denotes the weight vector for cell type *c* across *M* metacells. Then, the aggregated cell-type-level association 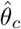 is given by:

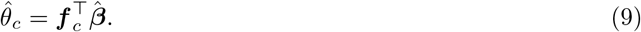

We estimate the variance of 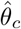 using a permutation-based null distribution. Specifically, we permute MAGMA z-score vector obtained by randomly shuffling the original gene-level MAGMA z-scores across genes, while keeping metacell specificity scores fixed. For each permutation *b* = 1, …, *B*, metacell-level association effects are recomputed using the same OLS framework described above, yielding null regression coefficients

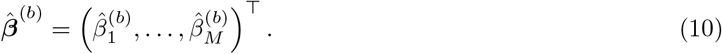

For each null metacell-level association estimate, corresponding significance weights are recalculated as

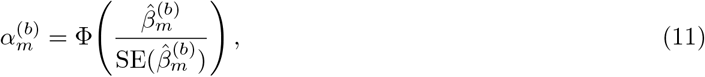

and the cell-type-specific normalized weight vector under permutation is defined as

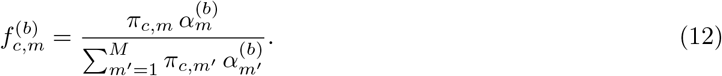

The corresponding null cell-type-level association estimate is then computed analogously to the observed statistic:

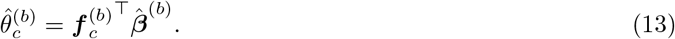

Because null metacell-level association estimates are approximately Gaussian, and the cell-type-level association statistic 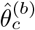 is computed as a weighted combination of these null metacell-level effects, the resulting null distribution of 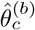 is also expected to approximately follow a Gaussian distribution. We therefore estimate the standard error of the observed cell-type-level association effect 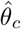 empirically from the permutation null distribution as

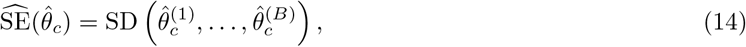

where SD(*·*) denotes the sample standard deviation across permutations. The final cell-type-level association z-score is then calculated as

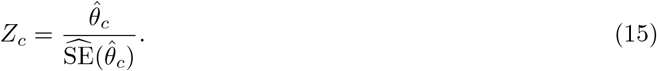

A one-sided p-value is computed as

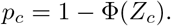

Notably, metacell-level association estimates are not independent because metacells can exhibit correlated gene expression patterns and transcriptional similarity. However, during permutation, only the disease-associated MAGMA z-scores are shuffled, while the metacell specificity matrix remains unchanged. Consequently, the correlation structure among metacells is preserved in both the observed and null statistics. This allows the empirical null distribution of 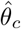 to implicitly account for metacell correlation and controls inflation of cell-type-level association statistics arising from dependent metacells.

#### Conducting heterogeneity-aware influential gene diagnostics

We adopted the idea from seismic [5], which uses DFBETAS [14] to prioritize influential genes that contribute to disease-cell type associations. However, because there can be substantial heterogeneity within a cell type, we modified this approach to prioritize influential genes that specifically contribute to disease associations from affected cell subpopulation.

For each metacell *m* and gene *i*, the *DFBETA*_*m,i*_, the unnormalized DFBETAS, is computed as the change in the metacell-level association coefficient after removing gene *i* from the regression model:

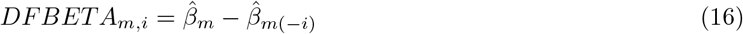

Then, for each gene i, the metacell-level influence statistics DFBETA_*m,i*_ is aggregated to the cell-type level using the same weight vector ***f***_*c*_ defined above. Let DFBETA_*·i*_ = (DFBETA_1,*i*_, …, DFBETA_*M,i*_)^*⊤*^ denote the vector of metacell-level DFBETA values for gene *i*. The cell-type-level DFBETA for gene *i* in cell type *c* is defined as:

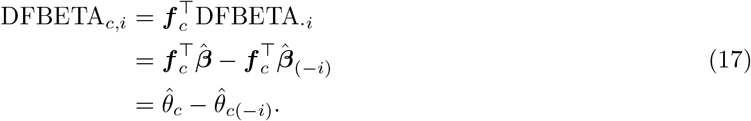

Theoretically, the cell-type-level DFBETAS statistic—defined as the standardized DFBETA—requires normalizing DFBETA_*c,i*_ by the standard error of the cell-type-level association after removing gene *i*, namely 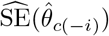. To compute this quantity exactly, we would need to re-estimate the metacell-level association significance without gene *i*, combined with cell type purity to get weight vector, and recompute the metacell-level covariance matrix after excluding gene *i*, finally propagate the updated covariance to the cell-type level to obtain 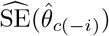. However, this procedure is computationally infeasible in practice. In our setting, the number of genes is large (e.g., >15,000 genes in the TM FACS dataset). Removing a single gene has a negligible impact on the metacell-level standard errors and, consequently, on the aggregated cell-type-level standard error. Therefore, the standardized statistic is approximated by using the full-model standard error 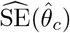, yielding:

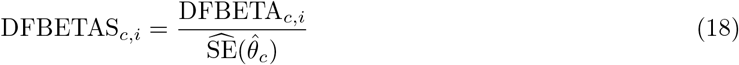

An empirical threshold of 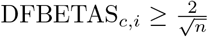 was used to identify genes contributing to disease-cell associations within affected cell populations, where *n* is the number of genes involved in association tests.

#### Accessing heterogeneous association within cell types

We quantified within-cell-type heterogeneity by measuring the proportion of individual cells belonging to metacells that are significantly associated within each cell type. To identify associated metacells, we applied a weighted Benjamini-Hochberg (BH) procedure [15]. This approach increases statistical power when prior information is available. In our setting, metacell purity is the prior information, as metacells from an associated cell type with higher cell-type purity are more likely to be associated.

Let *p*_*m*_ denote the p-value for metacell *m*. For each cell type *c*, let *ω*_*c,m*_ denote the corresponding weight assigned to metacell *m*, reflecting its purity with respect to cell type *c*. The weights are proportional to metacell purity and then normalized such that the total weight equals the number of metacells with non-zero purity for that cell type:

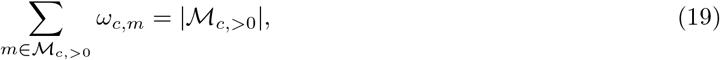

where ℳ_*c*,>0_ denotes the set of metacells with non-zero purity for cell type c. The weighted p-values are then defined as:

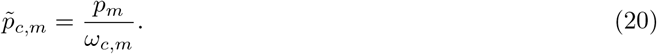

The weighted p-values are subsequently used in the standard BH procedure to control the false discovery rate (FDR) across metacells with non-zero purity for cell type *c*. This yields metacell-level FDR values FDR_*c,m*_ for each cell type *c* and metacell *m*, then each metacell-level FDR is mapped to its corresponding cells. Cells belonging to metacells with FDR_*c,m*_ < 0.1 are classified as significantly associated. To quantify within-cell-type heterogeneity, we computed the proportion of significantly associated cells within each cell type.

### Null and power analyses

#### Preparing simulation data

We randomly sampled 10,000 cells from the TM FACS dataset. Cell types represented by fewer than 10 cells were excluded, and genes expressed in fewer than 10 cells were removed. The resulting dataset was used as input for both the null and power analyses. Mouse gene expression was converted to human gene space using one-to-one orthologs obtained from NCBI Orthologs [81].

#### Conducting null simulation

In null simulations, we preserved the overall distribution of MAGMA z-scores while breaking any true association between cell type and disease by randomly shuffling MAGMA z-scores across genes. For each run, we randomly selected a disease from 84 traits, performed the shuffling, and compared the results against randomly sampled cells from the TM FACS dataset. Model calibration was assessed by comparing the distribution of calculated p-values to the expected uniform distribution under the null. We restricted this assessment to cell types with sufficient size, as small cell types yield unstable and statistically unreliable estimates; specifically, we evaluated calibration only for cell types with ≥ 40 cells.

#### Conducting power analysis

The goal of the power analysis was to evaluate whether methods can correctly identify an associated cell type when a true disease signal is present. In contrast to previous strategies used in scDRS and seismic, which simulate signal by perturbing gene expression profiles, we directly perturbed disease signals by synthesizing MAGMA gene-level z-scores. The limitations of simulation through gene expression perturbation are discussed in **Supplementary Note 5**.

We used the TM FACS dataset as the source for power simulations and preserve the original cell-type composition. In each simulation, one cell type is selected as the putative causal cell type. For each gene *g*, we compute the log fold change (logFC_*g*_) of its expression in the selected cell type compared to all other cell types. These log fold changes serve as the basis for simulated disease signals. A subset of genes is then designated as true causal genes. Gene selection probabilities determined by a softplus transformation of logFC_*g*_, which is further normalized to the sum of one. This design ensures that genes with higher logFC_*g*_ are more likely to be causal, while still allowing genes with near-zero or even negative logFC_*g*_ to be selected. The proportion of sampled causal genes is controlled by the parameter causal fraction. For causal genes, signal strength is modulated by multiplying logFC_*g*_ by a signal scaling factor, denoted as factor, and adding Gaussian noise *ε*_*g*_, where *ε*_*g*_ ~ *𝒩* (0, noise). The parameter noise controls the strength of the stochasticity. For non-causal genes, the signal arises purely from noise with the same variance as in casual genes. Formally, the simulated disease signal for each gene is defined as

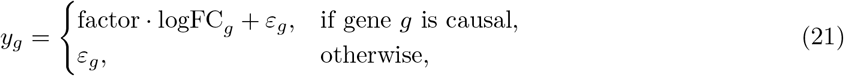

For each simulated disease signal, we matched its distribution to MAGMA gene-level z-scores. In each simulation run, we randomly selected a disease and mapped its MAGMA z-scores to genes by ranking *y*_*g*_ and assigning z-scores according to the same rank order, thereby preserving the empirical distribution.

In addition to controlling signal strength through causal fraction, factor, and noise, we introduced a setting to model heterogeneous disease signals within a cell type. Specifically, we constructed synthetic cell types composed of an associated subpopulation, defined as a localized patch of cells serving as the disease signal source, and a non-associated subpopulation sampled from a separate, but nearby region. A heterogeneity parameter *h ∈* [0, 1] controls the fraction of associated cells, with larger values of *h* corresponding to fewer associated cells, and smaller values (near 0) indicating that most cells in the cell type are associated.

To ensure biologically realistic sampling of continuous cell populations, we first clustered subsampled TM FACS data using Leiden clustering [82] at high resolution (resolution=5.0) to generate multiple localized cell patches. Associated cells were then sampled from clusters with sufficient size to meet the minimum required cell population size under a given heterogeneity level. For non-associated cells, rather than sampling arbitrarily distant populations, we ranked Leiden clusters by their proximity to the associated population using PCA-based Euclidean distance and sampled from clusters within a moderate distance range (30th to 50th percentile). To generate disease signals that emphasize the associated subpopulation, we computed log fold changes for both associated and non-associated groups and defined a contrast as

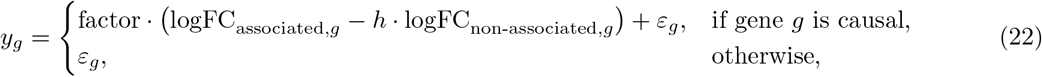

We introduce *h* to control the contribution of the non-associated signal, when *h*=0, the formulation reduces to Eq. 21, corresponding to the no-heterogeneity case. This contrast was transformed using an exponential function to further amplify association-specific signals, then normalized to sum to one and used as probabilities for selecting causal genes. The resulting signals were subsequently mapped to MAGMA z-scores.

To limit the number of simulation settings, we varied one parameter at a time while fixing all others at their default values. Specifically, the noise variance was set to noise = 1.0, the signal scaling factor to factor = 1.0, the fraction of causal genes to causal fraction = 0.01, and the heterogeneity level to heterogeneity level = 1.0. Under this design, we conducted four sets of experiments: (A) varying the fraction of causal genes (causal fraction ∈ 0.001, 0.005, 0.01, 0.05, 0.1) to assess sensitivity to signal sparsity; (B) varying the noise variance (noise ∈ 0.1, 0.5, 1.0, 2.0, 5.0) to evaluate robustness to noise; (C) varying the signal scaling factor (factor ∈ 0.1, 0.5, 1.0, 2.0, 5.0) to examine the effect of signal strength; and (D) varying the heterogeneity level (heterogeneity level ∈ 0.0, 0.2, 0.4, 0.6, 0.8) to model heterogeneous disease-cell type associations. For simulations based on whole cell types (varying noise, factor, and causal fraction), we restricted analysis to cell types with at least 40 cells. This constraint avoids instability in logFC_*g*_ estimates from small cell types, which can introduce noise-driven artifacts and artificially favor cell type-level methods such as seismic. When evaluating the effect of heterogeneity level, we further increased the minimum cell type size to 100 to ensure a sufficient number of associated cells under high heterogeneity. For each parameter setting, we generated 100 simulation runs and defined power as the proportion of runs in which the associated cell type was successfully detected (FDR ≤ 0.1). Each setting was repeated 10 times to assess variability in power.

#### Building disease-disease similarity graph

To group diseases based on metacell association profiles, we used metacell-level association z-scores as features and applied principal component analysis (PCA) to account for correlations among metacells and reduce redundancy in the feature space. Pairwise Euclidean distances between diseases were then computed in the PCA space, and a k-nearest neighbor (KNN) graph was constructed, then edge weights were subsequently smoothed using a Gaussian kernel. Disease clusters were identified using Louvain community detection optimization [83].

For comparison, we performed an analogous analysis using gene-level genetic risk profiles derived from MAGMA z-scores. To ensure comparability across traits, MAGMA z-scores were quantile-normalized. Euclidean distances were then calculated directly from the normalized gene-level profiles, followed by KNN graph construction and Louvain clustering using the same procedure as for metacell-level features.

We chose the resolution parameter at 2.5 to get more granular clusters that better distinguish broad categories.

#### Curating gene sets related to enterocyte functions and immune stress

We curated biologically relevant gene sets from the Molecular Signatures Database (MSigDB; Gene Ontology Biological Process collection) [84] to test whether UC-associated cells preferentially exhibit features of mature enterocyte function, whereas less-associated cells reflect reduced differentiation and enhanced resistance to immune stress. To capture canonical functions of differentiated enterocytes, we selected pathways related to electrolyte balance and fluid absorption [85, 86], including regulation of sodium ion transport (GO:0002028), chloride transport (GO:0006821), regulation of pH (GO:0006885), and water transport (GO:0006833). To represent cellular responses to immune stress, we included gene sets associated with endoplasmic reticulum (ER) stress signaling and apoptosis regulation [32, 87], specifically intrinsic apoptotic signaling pathway in response to endoplasmic reticulum stress (GO:0070059), negative regulation of endoplasmic reticulum stress-induced intrinsic apoptotic signaling pathway (GO:1902236), regulation of endoplasmic reticulum stress-induced intrinsic apoptotic signaling pathway (GO:1902235), and negative regulation of response to endoplasmic reticulum stress (GO:1903573). Broader ER stress terms that include positive regulation of ER stress were excluded to avoid capturing pathways reflecting stress exacerbation rather than resistance. We additionally incorporated gene sets related to epithelial maturation and maintenance, including intestinal epithelial cell differentiation (GO:0060575), intestinal epithelial structure maintenance (GO:0060729), and intestinal epithelial cell development (GO:0060576), to characterize differentiation status.

We pooled genes related to differentiated enterocyte function into a single gene set and combined genes associated with immune stress response and epithelial differentiation into a second set representing cellular homeostasis. After removing duplicates, these categories comprised 317 and 134 unique genes, respectively. To quantify relative program activity at the single-cell level, we computed the average log-normalized expression of each gene set per cell and derived a normalized expression fraction to reflect relative dominance between programs. Specifically, for each cell *i*, we calculated

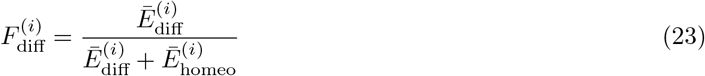

where 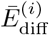 and 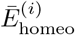 denote the average log-normalized expression of the differentiated-function and homeostasis gene sets for cell *i*, respectively. The corresponding cell homeostasis fraction was defined analogously.

#### Collecting and processing GWAS dataset

We collected GWAS summary statistics for 72 complex diseases from the set analyzed in the scDRS study [3]. These diseases span a broad range of categories, including immune, neurological, cardiovascular, liver-related, and lung-related disorders. In addition, we included nine additional traits to provide a more comprehensive representation of traits and phenotypes. Detailed information for all traits is provided in Supplementary Data 1. For autism spectrum disorder, we obtained GWAS summary statistics from Pedersen et al. [52].

GWAS summary statistics were aggregated to gene-level scores using MAGMA [2]. SNPs were mapped to genes using MAGMA’s default parameters, assigning SNPs within *±*10 kb of each gene to that gene.

#### Collecting and processing single cell data

We processed the TM FACS dataset [88]. The count matrix was downloaded directly from the published resource. No additional cell filtering was applied because the original dataset had already removed cells with fewer than 500 detected genes (genes with non-zero UMI counts). We further filtered genes by removing those expressed in fewer than 10 cells.

For the mouse colon dataset from Drokhlyansky et al. [49], the published data had already filtered cells with fewer than 1,000 detected genes. Therefore, we did not perform additional cell-level filtering. We removed genes expressed in fewer than 10 cells. For both datasets, we retained only genes with valid Entrez IDs. When multiple gene symbols mapped to the same Entrez ID, we kept the gene with the highest mean expression across cells. For visualization, we used the tSNE embedding provided in the original study rather than UMAP, as it showed clearer separation among cell groups.

For MetaQ input, raw count data are required. However, to compute metacell-specific expression scores, we need normalized counts. We first normalized counts by sequencing depth (scaled to 10,000 counts per cell), and then applied natural log transformation with a pseudocount of 1.

For mapping of gene from mouse to human, we used one-to-one orthologs from NCBI Orthologs [81].

#### Running benchmarked methods

For seismic [5], we did not apply the default filtering parameters. Seismic’s strict filtering removes a substantial number of genes, resulting in a marked decrease in disease-cell-type associations (**Supplementary Fig. 20**). For consistency across methods, we therefore did not filter any genes or cell types and discuss the problem of strict filtering in detail in **Supplementary Note 2**. Log-normalized expression data were used as input, together with the cross-species conversion map based on one-to-one orthologs from NCBI Orthologs[81]. Cell types with FDR ≤ 0.1 were considered significant.

For scDRS [3], we used default parameters (1,000 genes) with 1,000 Monte Carlo (MC) samples. The top 5% quantile of trait scores across cells within each cell type was used as the test statistic. Prior to running scDRS, TM FACS mouse expression data were converted to human gene space using one-to-one orthologs from NCBI Orthologs [81] to ensure consistent gene mapping across methods. Disease-associated gene sets were selected from top 1,000 genes in the intersection between converted expression genes and genes present in the MAGMA z-score list. Cell types with FDR ≤ 0.1 were considered significant.

Additional methodological details of seismic and scDRS are provided in **Supplementary Note 1**.

## Code and Data Availability

All code used for the analyses is available at https://github.com/krishnanlab/icepopanalysis. All processed single-cell data, GWAS summary statistics, and simulation data used in this study are available via Zenodo (https://doi.org/10.5281/zenodo.20146708). ICePop is released as a python package and is available at https://github.com/krishnanlab/icepop.

## Acknowledgement

We would like to thank all members of the Krishnan Lab for valuable discussions and feedback on the project.

## Funding

This work is supported by NIH R35 GM128765 and Simons Foundation 1017799 (to A.K.).

## Author contributions

H.Y. and A.K. designed the study. H.Y. developed the software. A.M. developed the interactive outputs.

K.S. performed code review and testing. H.Y. and K.S. performed the analyses. H.Y. interpreted the results.

J.G. provided feedback on the ASD analysis. H.Y. wrote the manuscript with feedback from A.M., K.S., J.G., and A.K.

## Notes

https://doi.org/10.5281/zenodo.20146708

### Competing Interest Statement

The authors have declared no competing interest.

### Summary of Updates

In methods, we added additional filtering on trivial disease heterogeneity. We reran the whole pipeline, thus all figures except fig. 1 were updated. Supplemental files updated.

## References

[1] Visscher, P. M. et al. 10 years of gwas discovery: biology, function, and translation. The American journal of human genetics 101, 5–22 (2017).

[2] De Leeuw, C. A., Mooij, J. M., Heskes, T. & Posthuma, D. Magma: generalized gene-set analysis of gwas data. PLoS computational biology 11, e1004219 (2015).

[3] Zhang, M. J. et al. Polygenic enrichment distinguishes disease associations of individual cells in single-cell rna-seq data. Nature genetics 54, 1572–1580 (2022).

[4] Boyle, E. A., Li, Y. I. & Pritchard, J. K. An expanded view of complex traits: from polygenic to omnigenic. Cell 169, 1177–1186 (2017).

[5] Lai, Q., Dannenfelser, R.Roussarie, J.-P. & Yao, V. Disentangling associations between complex traits and cell types with seismic. bioRxiv (2024).

[6] Fan, J. et al. Characterizing transcriptional heterogeneity through pathway and gene set overdispersion analysis. Nature methods 13, 241–244 (2016).

[7] Cembrowski, M. S. & Spruston, N. Heterogeneity within classical cell types is the rule: lessons from hippocampal pyramidal neurons. Nature Reviews Neuroscience 20, 193–204 (2019).

[8] Keren-Shaul, H. et al. A unique microglia type associated with restricting development of alzheimer’s disease. Cell 169, 1276–1290 (2017).

[9] Friedman, B. A. et al. Diverse brain myeloid expression profiles reveal distinct microglial activation states and aspects of alzheimer’s disease not evident in mouse models. Cell reports 22, 832–847 (2018).

[10] Mathys, H. et al. Single-cell transcriptomic analysis of alzheimer’s disease. Nature 570, 332–337 (2019).

[11] Bilous, M., Hérault, L., Gabriel, A. A., Teleman, M. & Gfeller, D. Building and analyzing metacells in single-cell genomics data. Molecular Systems Biology 20, 744 (2024).

[12] A single-cell transcriptomic atlas characterizes ageing tissues in the mouse. Nature 583, 590–595 (2020).

[13] Li, Y. et al. Metaq: fast, scalable and accurate metacell inference via single-cell quantization. Nature Communications 16, 1205 (2025).

[14] Belsley, D. A., Kuh, E. & Welsch, R. E. Regression diagnostics: Identifying influential data and sources of collinearity (John Wiley & Sons, 2005).

[15] Genovese, C. R., Roeder, K. & Wasserman, L. False discovery control with p-value weighting. Biometrika 93, 509–524 (2006).

[16] Watanabe, K., Taskesen, E., Van Bochoven, A. & Posthuma, D. Functional mapping and annotation of genetic associations with fuma. Nature communications 8, 1826 (2017).

[17] de Leeuw, C. A., Stringer, S., Dekkers, I. A., Heskes, T. & Posthuma, D. Conditional and interaction gene-set analysis reveals novel functional pathways for blood pressure. Nature communications 9, 3768 (2018).

[18] Makhmutova, M. et al. Pancreatic β-cells communicate with vagal sensory neurons. Gastroenterology 160, 875–888 (2021).

[19] Slominski, A. et al. Hair follicle pigmentation. Journal of Investigative Dermatology 124, 13–21 (2005).

[20] Saleh, M. et al. Diastolic blood pressure predicts coronary plaque volume in patients with coronary artery disease. Atherosclerosis 277, 34–41 (2018).

[21] Yan, H. et al. Peptide-sirna nanoparticles targeting nf-κb p50 mitigate experimental abdominal aortic aneurysm progression and rupture. Biomaterials advances 139, 213009 (2022).

[22] Travers, J. G., Kamal, F. A., Robbins, J., Yutzey, K. E. & Blaxall, B. C. Cardiac fibrosis: the fibroblast awakens. Circulation research 118, 1021–1040 (2016).

[23] Quansah, E. et al. Intestinal epithelial barrier integrity investigated by label-free techniques in ulcerative colitis patients. Scientific Reports 13, 2681 (2023).

[24] Heller, F. et al. Interleukin-13 is the key effector th2 cytokine in ulcerative colitis that affects epithelial tight junctions, apoptosis, and cell restitution. Gastroenterology 129, 550–564 (2005).

[25] Neurath, M. F. Strategies for targeting cytokines in inflammatory bowel disease. Nature Reviews Immunology 24, 559–576 (2024).

[26] Kiesslich, R. et al. Local barrier dysfunction identified by confocal laser endomicroscopy predicts relapse in inflammatory bowel disease. Gut 61, 1146–1153 (2012).

[27] Villanacci, V., Del Sordo, R., Parigi, T. L., Leoncini, G. & Bassotti, G. Inflammatory bowel diseases: does one histological score fit all? Diagnostics 13, 2112 (2023).

[28] Braga, V. Spatial integration of e-cadherin adhesion, signalling and the epithelial cytoskeleton. Current opinion in cell biology 42, 138–145 (2016).

[29] Suenaga, M. et al. Role of enterocyte-specific gene polymorphisms in response to adjuvant treatment for stage iii colorectal cancer. Pharmacogenetics and genomics 31, 10–16 (2021).

[30] Chan, C. W. et al. Gastrointestinal differentiation marker cytokeratin 20 is regulated by homeobox gene cdx1. Proceedings of the National Academy of Sciences 106, 1936–1941 (2009).

[31] Piguet, P. F., Vesin, C., Guo, J., Donati, Y. & Barazzone, C. Tnf-induced enterocyte apoptosis in mice is mediated by the tnf receptor 1 and does not require p53. European journal of immunology 28, 3499–3505 (1998).

[32] Luo, K. & Cao, S. S. Endoplasmic reticulum stress in intestinal epithelial cell function and inflammatory bowel disease. Gastroenterology research and practice 2015, 328791 (2015).

[33] Hayashi, H., Suruga, K. & Yamashita, Y. Regulation of intestinal cl-/hco3-exchanger slc26a3 by intracellular ph. American Journal of Physiology-Cell Physiology 296, C1279–C1290 (2009).

[34] Rocafull, M. A., Thomas, L. E., Barrera, G. J. & Del Castillo, J. R. Differential expression of p-type atpases in intestinal epithelial cells: identification of putative new atp1a1 splice-variant. Biochemical and biophysical research communications 391, 152–158 (2010).

[35] Escudero-Hernández, C., Münch, A., Østvik, A.-E., Granlund, A. v. B. & Koch, S. The water channel aquaporin 8 is a critical regulator of intestinal fluid homeostasis in collagenous colitis. Journal of Crohn’s and Colitis 14, 962–973 (2020).

[36] Wang, R. et al. Type 2 cgmp-dependent protein kinase regulates proliferation and differentiation in the colonic mucosa. American Journal of Physiology-Gastrointestinal and Liver Physiology 303, G209–G219 (2012).

[37] Wang, Y. et al. Mitochondrial protein if1 is a potential regulator of glucagon-like peptide (glp-1) secretion function of the mouse intestine. Acta Pharmaceutica Sinica B 11, 1568–1577 (2021).

[38] Fujimoto, K. et al. Regulation of intestinal homeostasis by the ulcerative colitis-associated gene rnf186. Mucosal immunology 10, 446–459 (2017).

[39] Raslan, A. A. et al. Lung injury-induced activated endothelial cell states persist in aging-associated progressive fibrosis. Nature communications 15, 5449 (2024).

[40] Wakim, L. M., Gupta, N., Mintern, J. D. & Villadangos, J. A. Enhanced survival of lung tissue-resident memory cd8+ t cells during infection with influenza virus due to selective expression of ifitm3. Nature immunology 14, 238–245 (2013).

[41] Stewart, W. G., Hejl, C. D., Guleria, R. S. & Gupta, S. Thymosin β4 stabilizes hypoxia induced brain microvascular endothelial cell dysfunction through s1pr1 dependent mechanisms. Scientific Reports (2025).

[42] Caporarello, N. et al. Dysfunctional erg signaling drives pulmonary vascular aging and persistent fibrosis. Nature communications 13, 4170 (2022).

[43] Watanabe, K. et al. A global overview of pleiotropy and genetic architecture in complex traits. Nature genetics 51, 1339–1348 (2019).

[44] An, S. Y. et al. Keratin-mediated hair growth and its underlying biological mechanism. Communications Biology 5, 1270 (2022).

[45] Balañá, M. E., Charreau, H. E. & Leirós, G. J. Epidermal stem cells and skin tissue engineering in hair follicle regeneration. World journal of stem cells 7, 711 (2015).

[46] Zhou, S. et al. Crosstalk between endothelial cells and dermal papilla entails hair regeneration and angiogenesis during aging. Journal of Advanced Research 70, 339–353 (2025).

[47] McElhanon, B. O., McCracken, C., Karpen, S. & Sharp, W. G. Gastrointestinal symptoms in autism spectrum disorder: a meta-analysis. Pediatrics 133, 872–883 (2014).

[48] Sotelo-Orozco, J. & Hertz-Picciotto, I. The association between gastrointestinal issues and psychometric scores in children with autism spectrum disorder, developmental delays, down syndrome, and typical development. Journal of Autism and Developmental Disorders 55, 2452–2462 (2025).

[49] Drokhlyansky, E. et al. The human and mouse enteric nervous system at single-cell resolution. Cell 182, 1606–1622 (2020).

[50] Chatterjee, I. et al. Chd8 regulates gut epithelial cell function and affects autism-related behaviors through the gut-brain axis. Translational Psychiatry 13, 305 (2023).

[51] Ji, P. et al. Single-cell delineation of the microbiota-gut-brain axis: Probiotic intervention in chd8 haploinsufficient mice. Cell Genomics 5 (2025).

[52] Pedersen, E. M. et al. Adult: An efficient and robust time-to-event gwas. Nature communications 14, 5553 (2023).

[53] Sun, N., Cao, L.-S., Xia, W.-Y.Wang, J.-M. & Wu, Q.-F. Gut sensory neurons as regulators of neuro-immune-microbial interactions: from molecular mechanisms to precision therapy for ibd/ibs. Journal of Neuroinflammation 22, 172 (2025).

[54] Wang, X. et al. The enteric nervous system deficits in autism spectrum disorder. Frontiers in neuroscience 17, 1101071 (2023).

[55] Hj, C. Enteric neural regulation of mucosal secretion. Physiology of the gastrointestinal tract 737–762 (2006).

[56] Tingler, A. M. et al. Children with autism spectrum disorder and chronic gastrointestinal symptoms have alterations in intestinal neurotransmitter pathways. Digestive Diseases and Sciences 1–19 (2026).

[57] Gomez-Frittelli, J. et al. Synaptic cell adhesion molecule cdh6 identifies a class of sensory neurons with novel functions in colonic motility. Elife 13, RP101043 (2025).

[58] Sharkey, K. A. & Mawe, G. M. The enteric nervous system. Physiological reviews 103, 1487–1564 (2023).

[59] Furness, J. B., Jones, C., Nurgali, K. & Clerc, N. Intrinsic primary afferent neurons and nerve circuits within the intestine. Progress in neurobiology 72, 143–164 (2004).

[60] Xie, Z. et al. Enteric neuronal piezo1 maintains mechanical and immunological homeostasis by sensing force. Cell 188, 2417–2432 (2025).

[61] Banerjee-Basu, S. & Packer, A. Sfari gene: an evolving database for the autism research community (2010).

[62] McCluskey, K. et al. Autism gene variants disrupt enteric neuron migration and cause gastrointestinal dysmotility. biorxiv. Preprint 2024 (2024).

[63] Robinson, B. G., Oster, B. A., Robertson, K. & Kaltschmidt, J. A. Loss of asd-related molecule cntnap2 affects colonic motility in mice. Frontiers in Neuroscience 17, 1287057 (2023).

[64] Anderson, B. et al. Examination of association of genes in the serotonin system to autism. Neurogenetics 10, 209–216 (2009).

[65] Bilous, M. et al. Metacells untangle large and complex single-cell transcriptome networks. BMC bioinformatics 23, 336 (2022).

[66] Persad, S. et al. Seacells infers transcriptional and epigenomic cellular states from single-cell genomics data. Nature biotechnology 41, 1746–1757 (2023).

[67] Smillie, C. S. et al. Intra-and inter-cellular rewiring of the human colon during ulcerative colitis. Cell 178, 714–730 (2019).

[68] Gao, H. et al. Endoplasmic reticulum stress of gut enterocyte and intestinal diseases. Frontiers in Molecular Biosciences 9, 817392 (2022).

[69] Qiu, W. et al. Puma-mediated intestinal epithelial apoptosis contributes to ulcerative colitis in humans and mice. The Journal of clinical investigation 121, 1722–1732 (2011).

[70] Iwamoto, M., Koji, T., Makiyama, K., Kobayashi, N. & Nakane, P. K. Apoptosis of crypt epithelial cells in ulcerative colitis. The Journal of pathology 180, 152–159 (1996).

[71] Fung, C. & Vanden Berghe, P. Functional circuits and signal processing in the enteric nervous system. Cellular and Molecular Life Sciences 77, 4505–4522 (2020).

[72] Said, H. M. Physiology of the gastrointestinal tract (Academic Press, 2018).

[73] Laurenti, E. & Göttgens, B. From haematopoietic stem cells to complex differentiation landscapes. Nature 553, 418–426 (2018).

[74] Ben-Kiki, O., Bercovich, A., Lifshitz, A. & Tanay, A. Metacell-2: a divide-and-conquer metacell algorithm for scalable scrna-seq analysis. Genome Biology 23, 100 (2022).

[75] Timshel, P. N., Thompson, J. J. & Pers, T. H. Genetic mapping of etiologic brain cell types for obesity. Elife 9, e55851 (2020).

[76] Jagadeesh, K. A. et al. Identifying disease-critical cell types and cellular processes by integrating single-cell rna-sequencing and human genetics. Nature genetics 54, 1479–1492 (2022).

[77] Pividori, M. et al. Projecting genetic associations through gene expression patterns highlights disease etiology and drug mechanisms. Nature communications 14, 5562 (2023).

[78] Szklarczyk, D. et al. The string database in 2025: protein networks with directionality of regulation. Nucleic acids research 53, D730–D737 (2025).

[79] Su, C. et al. Cell-type-specific co-expression inference from single cell rna-sequencing data. Nature Communications 14, 4846 (2023).

[80] Aibar, S. et al. Scenic: single-cell regulatory network inference and clustering. Nature methods 14, 1083–1086 (2017).

[81] Oh, D.-H. et al. Ncbi orthologs: public resource and scalable method for computing high-precision orthologs across eukaryotic genomes. Journal of Molecular Evolution 93, 843–859 (2025).

[82] Traag, V. A., Waltman, L. & Van Eck, N. J. From louvain to leiden: guaranteeing well-connected communities. Scientific reports 9, 5233 (2019).

[83] Blondel, V. D., Guillaume, J.-L., Lambiotte, R. & Lefebvre, E. Fast unfolding of communities in large networks. Journal of statistical mechanics: theory and experiment 2008, P10008 (2008).

[84] Liberzon, A. et al. The molecular signatures database hallmark gene set collection. Cell systems 1, 417–425 (2015).

[85] Priyamvada, S. et al. Mechanisms underlying dysregulation of electrolyte absorption in inflammatory bowel disease–associated diarrhea. Inflammatory bowel diseases 21, 2926–2935 (2015).

[86] Barrett, K. E. & Keely, S. J. Electrolyte secretion and absorption in the small intestine and colon. Yamada’s Textbook of Gastroenterology 283–312 (2022).

[87] Ma, X. et al. Intestinal epithelial cell endoplasmic reticulum stress and inflammatory bowel disease pathogenesis: an update review. Frontiers in immunology 8, 1271 (2017).

[88] Schaum, N. et al. Single-cell transcriptomics of 20 mouse organs creates a tabula muris: The tabula muris consortium. Nature 562, 367 (2018).

