## Supplementary material for "Linking Genetic Risk to Disease-Relevant Cellular States via Metacell-Informed Modeling with ICePop"

### Supplementary Figures

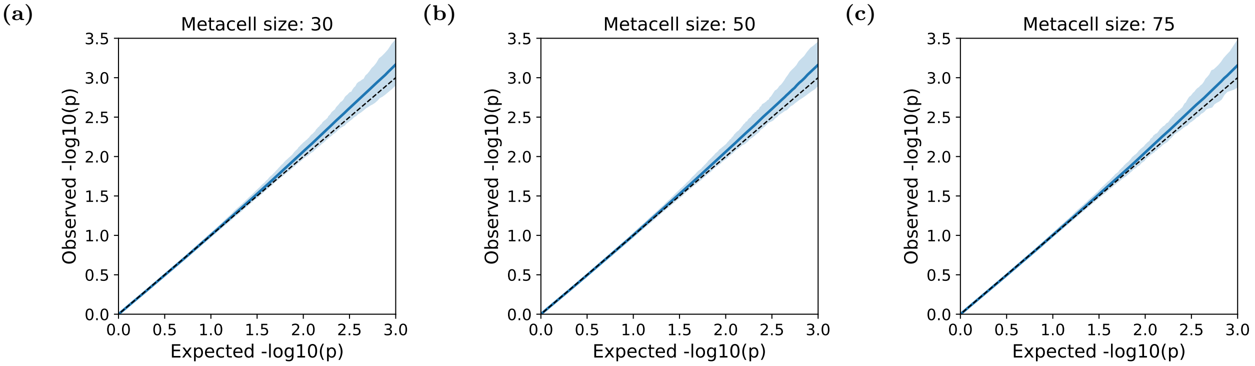


**Figure S1: Quantile-quantile (QQ) plots for metacell-level null simulations across different metacell sizes.** Null distributions were generated by permuting MAGMA z-scores across 10,000 runs. Associations were computed at the metacell level without cell-type aggregation. For each metacell, p-values were compared to the expected null distribution to construct QQ curves. The x-axis shows expected p-values under the null, and the y-axis shows observed p-values derived from associations between permuted MAGMA z-scores and subsets of 10,000 cells sampled from the Tabula Muris FACS dataset. The shaded region represents variability of QQ curves (5th-95th percentile range). The dashed line indicates the theoretical null expectation, and the solid line shows the mean observed $-\log_{10}(p)$. Panels a-c illustrate variability across individual metacells for metacell sizes of 30, 50, and 75, respectively.

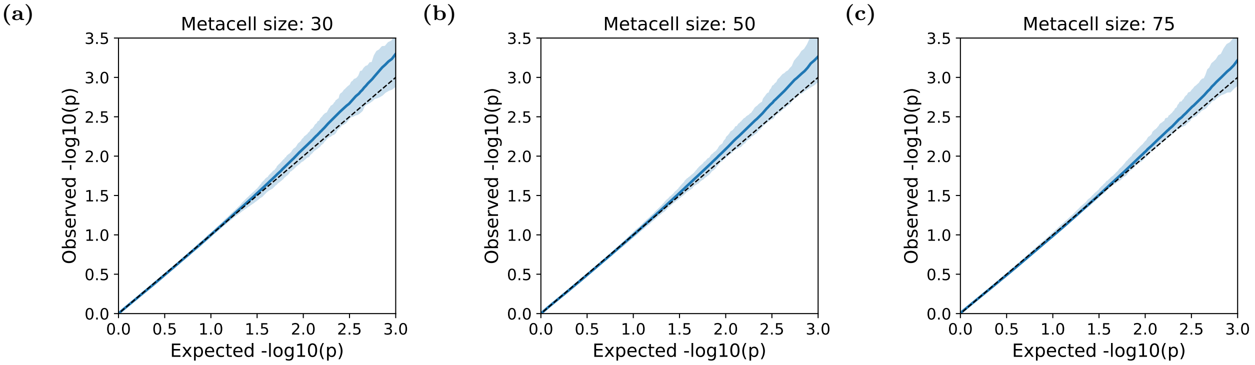


**Figure S2: Quantile-quantile (QQ) plots for cell-type-level null simulations across different metacell sizes.** Null distributions were generated by permuting MAGMA z-scores over 10,000 runs. Metacell-level associations were first computed and then aggregated to the cell-type level prior to significance testing. For each cell type, aggregated p-values were compared to the expected null distribution to construct QQ curves. The x-axis shows expected p-values under the null, and the y-axis shows observed p-values obtained from associations between permuted MAGMA z-scores and subsets of 10,000 cells sampled from the Tabula Muris FACS dataset. The shaded region represents variability of QQ curves (5th-95th percentile range). The dashed line indicates the theoretical null expectation, and the solid line shows the mean observed $-\log_{10}(p)$ across runs. Panels a-c show variability across cell types for disease-cell-type associations aggregated from metacells of sizes 30, 50, and 75, respectively.

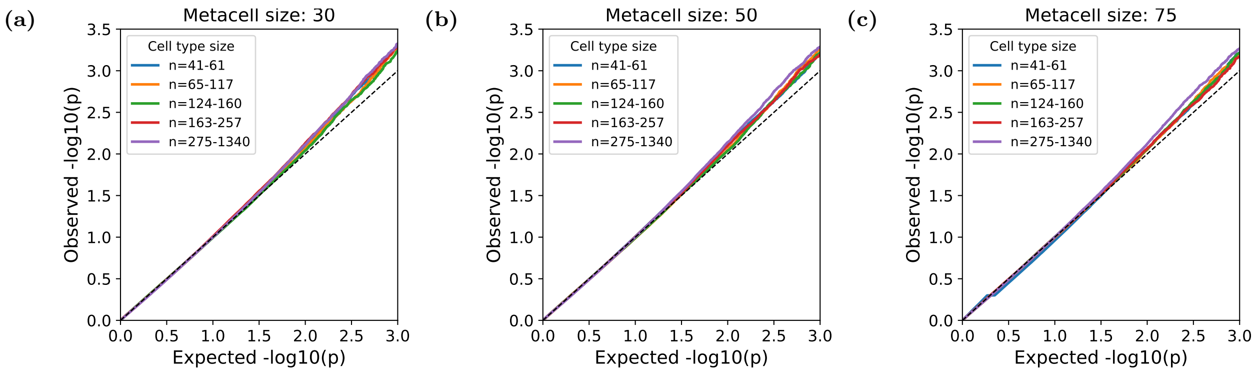


**Figure S3: Quantile-quantile (QQ) plots for aggregated cell-type-level null simulations, stratified by cell-type size.** Null distributions were generated by permuting MAGMA z-scores over 10,000 runs. Metacell-level associations were first computed and then aggregated to the cell-type level prior to significance testing. The x-axis shows expected p-values under the null, and the y-axis shows observed p-values from disease-cell-type association tests derived from these aggregated cell-type statistics using subsets of 10,000 cells sampled from the Tabula Muris FACS dataset. The dashed line indicates the theoretical null expectation. Solid lines represent the mean observed $-\log_{10}(p)$ across cell types, with colors corresponding to five cell-type-size bins. Panels a-c display averaged QQ curves for aggregated disease-cell-type associations from metacell sizes of 30, 50, and 75, respectively.

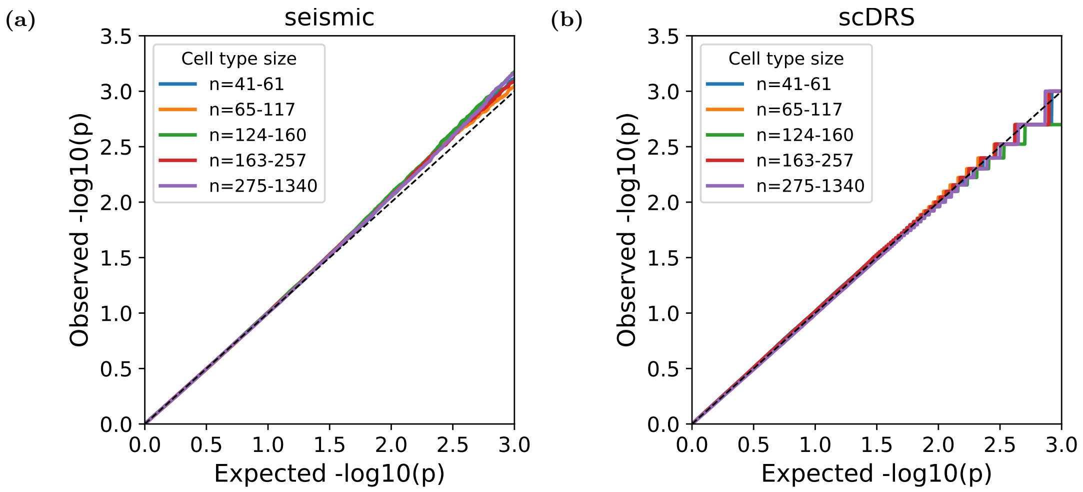


**Figure S4: Quantile-quantile (QQ) plots for cell-type-level null simulations for seismic and scDRS, stratified by cell-type size.** Null distributions were generated by permuting MAGMA z-scores over 10,000 runs. The x-axis shows expected p-values under the null, and the y-axis shows observed p-values from disease-cell-type association tests computed using permuted MAGMA z-scores and subsets of 10,000 cells sampled from the Tabula Muris FACS dataset. The dashed line indicates the theoretical null expectation, and solid lines represent the mean observed $-\log_{10}(p)$ across cell types. In panels a and b, colored lines represent averaged QQ curves for different cell-type-size bins, with panel a corresponding to seismic and panel b to scDRS.

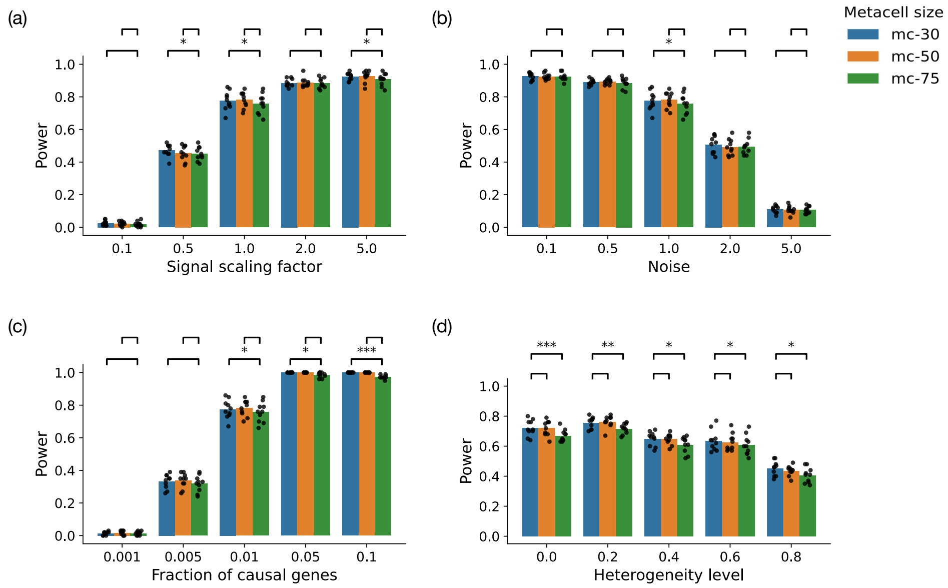


**Figure S5: Power comparisons for disease-cell-type association across metacell sizes under four simulation parameters.** MAGMA z-scores were synthetically generated from cell-type-specific expression profiles, with signal strength controlled by four parameters: fraction of causal genes (a), noise (b), signal scaling factor (c), and heterogeneity level (d). Each bar represents power which is calculated as the proportion of simulations (out of 100 independent runs) in which the causal cell type was correctly identified as significant at an FDR threshold of 0.1. Black dots indicate variability across 10 repeated runs under the same parameter setting. Analyses were performed on a subset of 10,000 cells from the Tabula Muris FACS dataset. Blue, orange, and green bars correspond to disease associations aggregated from metacells of sizes 30, 50, and 75, respectively. Statistical significance for pairwise power differences between metacell size 75 and sizes 50 or 30 is denoted by * ($p < 0.05$), ** ($p < 0.01$), and *** ($p < 0.001$). In each panel, one parameter is varied while the remaining three are held constant. Unless otherwise specified, default parameter values are: fraction of causal genes = 0.01, noise = 1.0, signal scaling factor = 1.0, and heterogeneity level = 0.0.

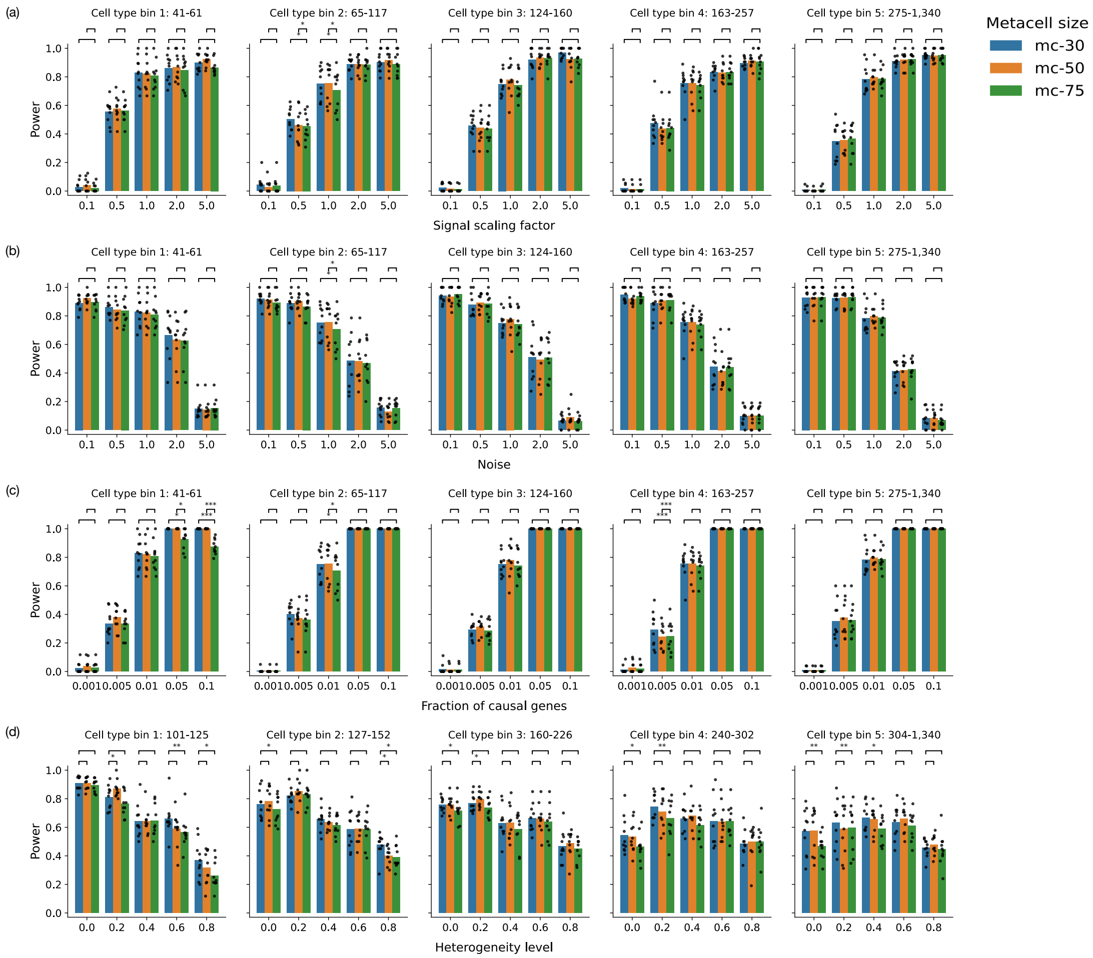


**Figure S6: Power comparisons for disease-cell-type associations across metacell sizes under four simulation parameters, stratified by cell-type size.** MAGMA z-scores were synthetically generated from cell-type-specific expression profiles, with signal strength controlled by four parameters: fraction of causal genes (a), noise (b), signal scaling factor (c), and heterogeneity level (d). Each bar represents power which is calculated as the proportion of simulations (out of 100 independent runs) in which the causal cell type was correctly identified as significant at an FDR threshold of 0.1. Black dots indicate variability across 10 repeated runs under the same parameter setting. Analyses were performed on a subset of 10,000 cells from the Tabula Muris FACS dataset. Blue, orange, and green bars correspond to disease associations aggregated from metacells of sizes 30, 50, and 75, respectively. Power are stratified across five cell-type-size bins. Statistical significance for pairwise power differences between metacell size 75 and sizes 50 or 30 is denoted by * ($p < 0.05$), ** ($p < 0.01$), and *** ($p < 0.001$). In each panel, one parameter is varied while the remaining three are held constant. Unless otherwise specified, default parameter values are: fraction of causal genes = 0.01, noise = 1.0, signal scaling factor = 1.0, and heterogeneity level = 0.0.

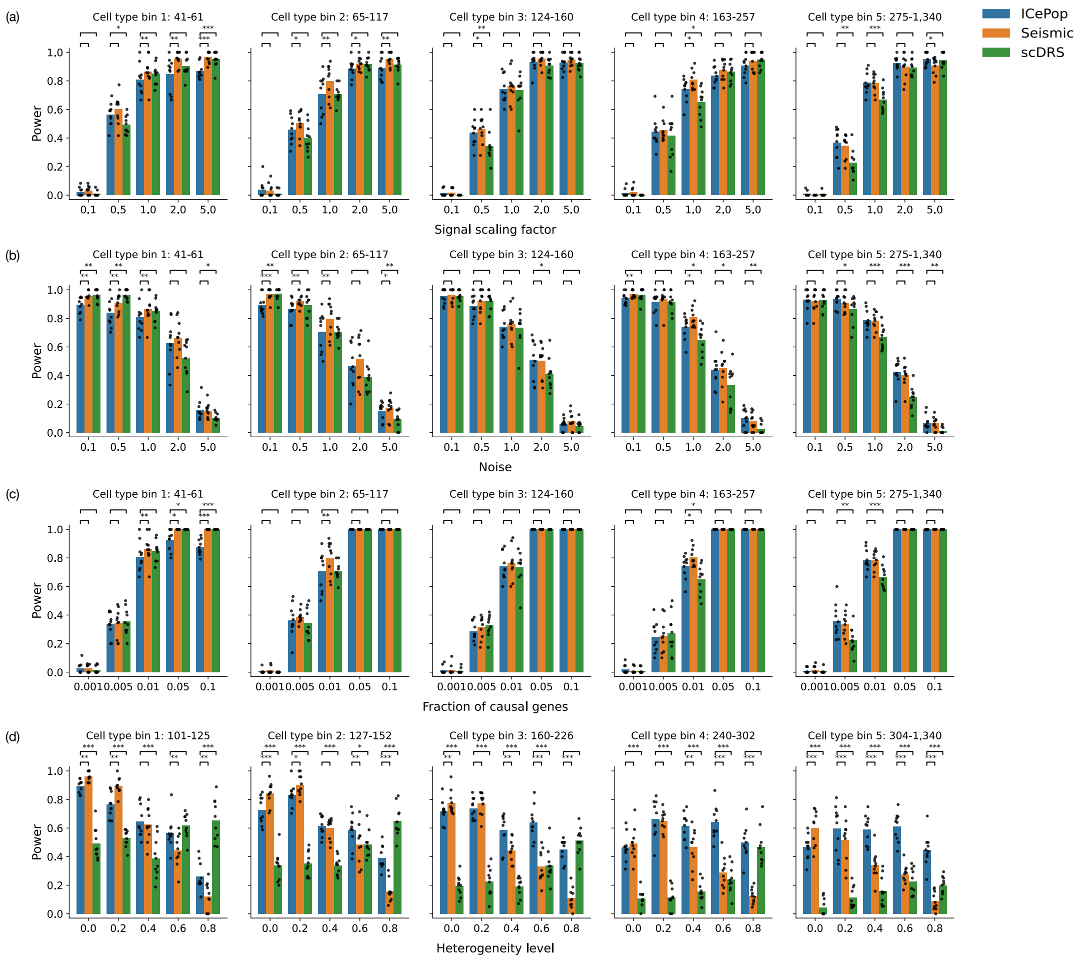


**Figure S7: Power comparisons for disease-cell-type associations across ICePop, seismic, and scDRS under four simulation parameters, stratified by cell-type size.** MAGMA z-scores were synthetically generated from cell-type-specific expression profiles, with signal strength controlled by four parameters: fraction of causal genes (a), noise level (b), signal scaling factor (c), and heterogeneity level (d). Each bar represents power which is calculated as the proportion of simulations (out of 100 independent runs) in which the causal cell type was correctly identified as significant at an FDR threshold of 0.1. Black dots indicate power variability across 10 repeated runs under the same parameter setting. Analyses were performed on a subset of 10,000 cells from the Tabula Muris FACS dataset. Blue, orange, and green bars correspond to disease-cell-type associations identified by ICePop (aggregated from metacells of size 75), seismic, and scDRS, respectively. Power estimates are stratified across five cell-type-size bins. Statistical significance for pairwise power differences between ICePop and seismic or scDRS is denoted by * ($p < 0.05$), ** ($p < 0.01$), and *** ($p < 0.001$). In each panel, one parameter is varied while the remaining three are held constant. Unless otherwise specified, default parameter values are: fraction of causal genes = 0.01, noise = 1.0, signal scaling factor = 1.0, and heterogeneity level = 0.0.

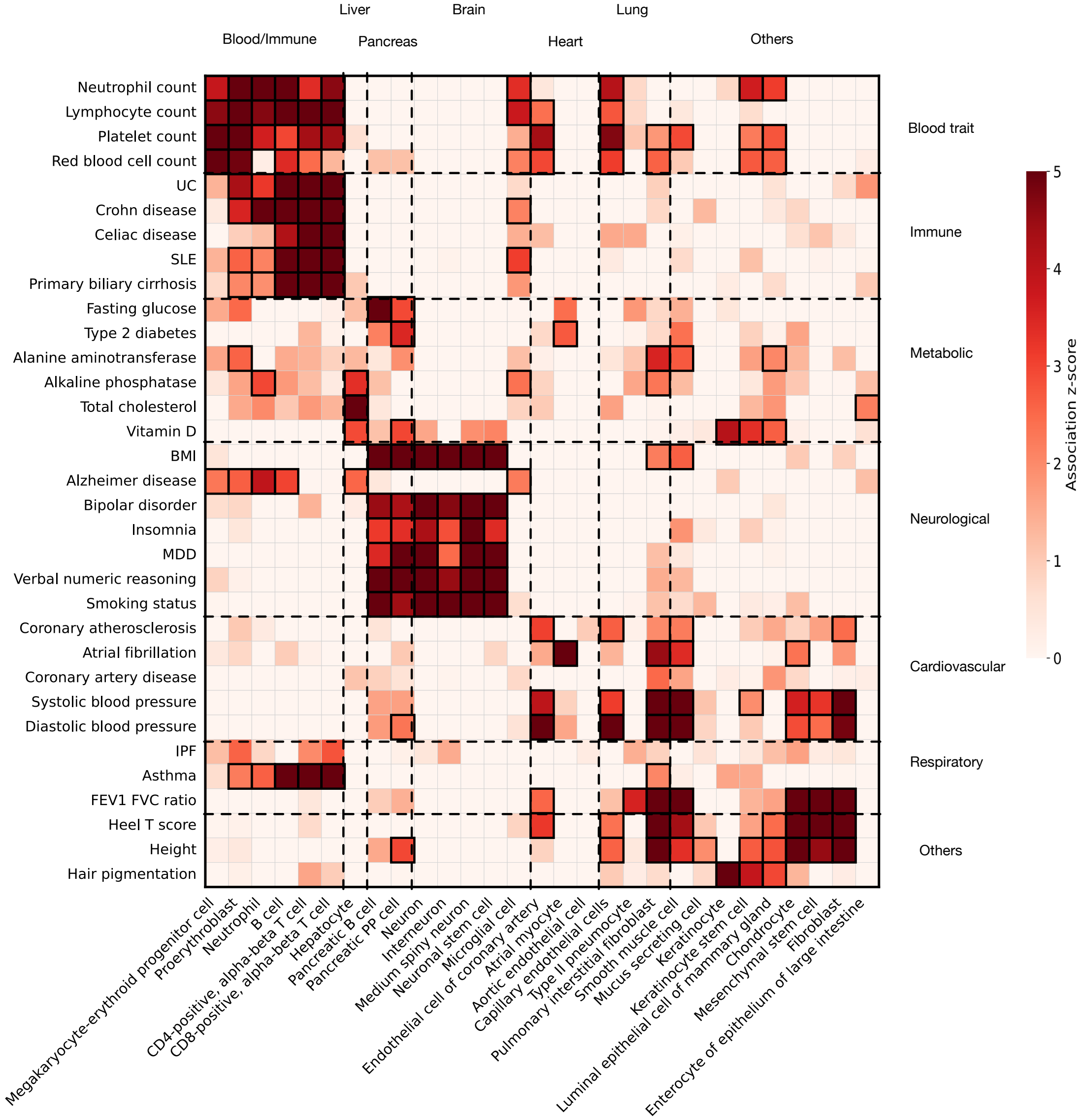


**Figure S8: Heatmap of seismic disease-cell-type association results for 33 representative traits and 29 cell types from the Tabula Muris FACS dataset.** Rows correspond to diseases or traits, and columns correspond to cell types, grouped into seven disease categories and seven tissues or organs. Colors indicate cell-type association z-scores from seismic, with negative values truncated to zero. Black rectangles denote significant associations, and cross symbols indicate heterogeneity of disease association, defined as significant association (FDR $\leq 0.1$) present in more than 20% but fewer than 80% of cells within a cell type. Trait abbreviations include ulcerative colitis (UC), systemic lupus erythematosus (SLE), body mass index (BMI), major depressive disorder (MDD), and idiopathic pulmonary fibrosis (IPF). Association results for all 81 traits and 120 cell types for seismic are provided in Supplementary Data 2.

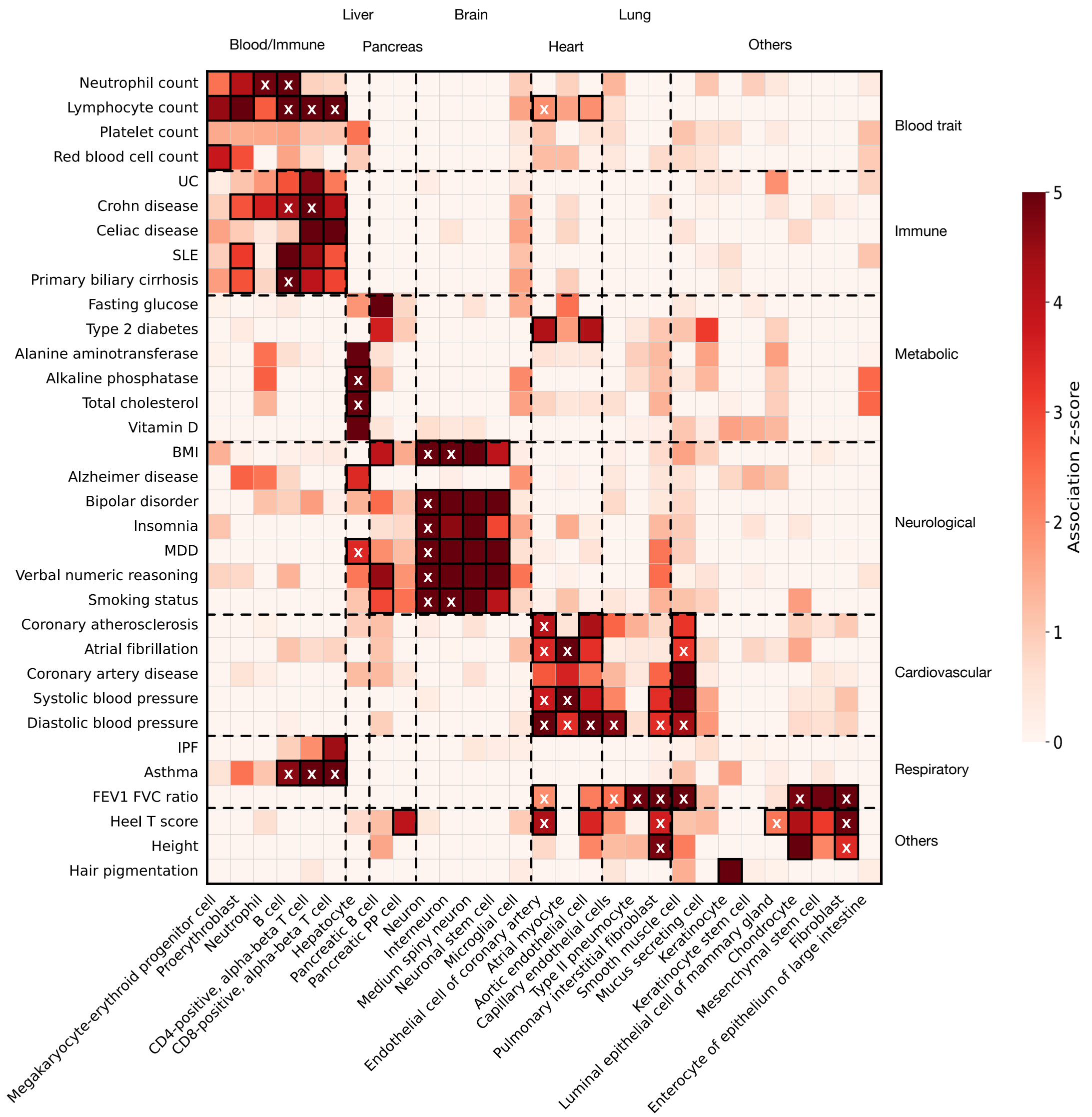


**Figure S9: Heatmap of scDRS disease-cell-type association results for 33 representative traits and 29 cell types from the Tabula Muris FACS dataset.** Rows correspond to diseases or traits, and columns correspond to cell types, grouped into seven disease categories and seven tissues or organs. Colors indicate cell-type association z-scores from scDRS, with negative values truncated to zero. Black rectangles denote significant associations, and cross symbols indicate heterogeneity of disease association, defined as significant association (FDR $\leq 0.1$) present in more than 20% but fewer than 80% of cells within a cell type. Trait abbreviations include ulcerative colitis (UC), systemic lupus erythematosus (SLE), body mass index (BMI), major depressive disorder (MDD), and idiopathic pulmonary fibrosis (IPF). Association results for all 81 traits and 120 cell types for scDRS are provided in Supplementary Data 1.

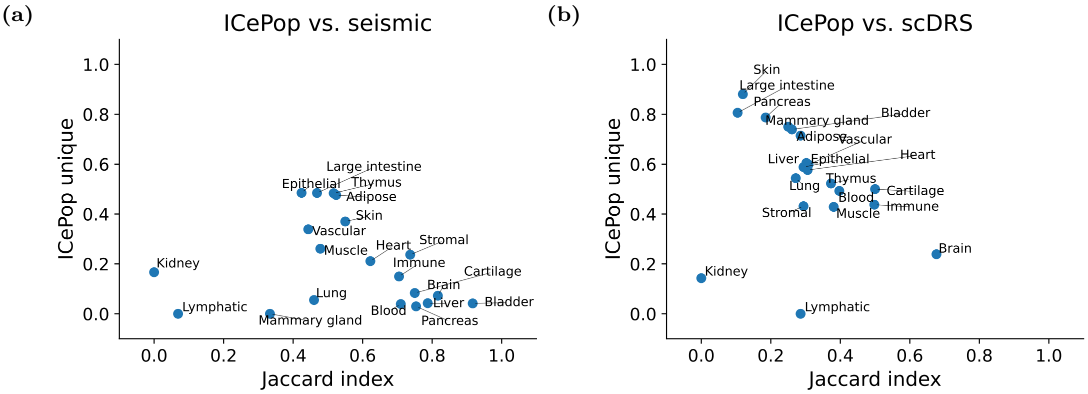


**Figure S10: Comparison of disease-cell-type associations between ICePop and seismic or scDRS across tissues.** Scatter plots showing the shared and unique disease-cell-type identifications between ICePop and seismic (a) or scDRS (b). Each point represents a tissue from the Tabula Muris FACS dataset. In both plots, the x-axis shows the Jaccard index, indicating the proportion of shared identifications between methods. The y-axis represents the fraction of ICePop-specific identifications, calculated as the number of associations uniquely identified by ICePop divided by the union of associations identified by ICePop and seismic or scDRS.

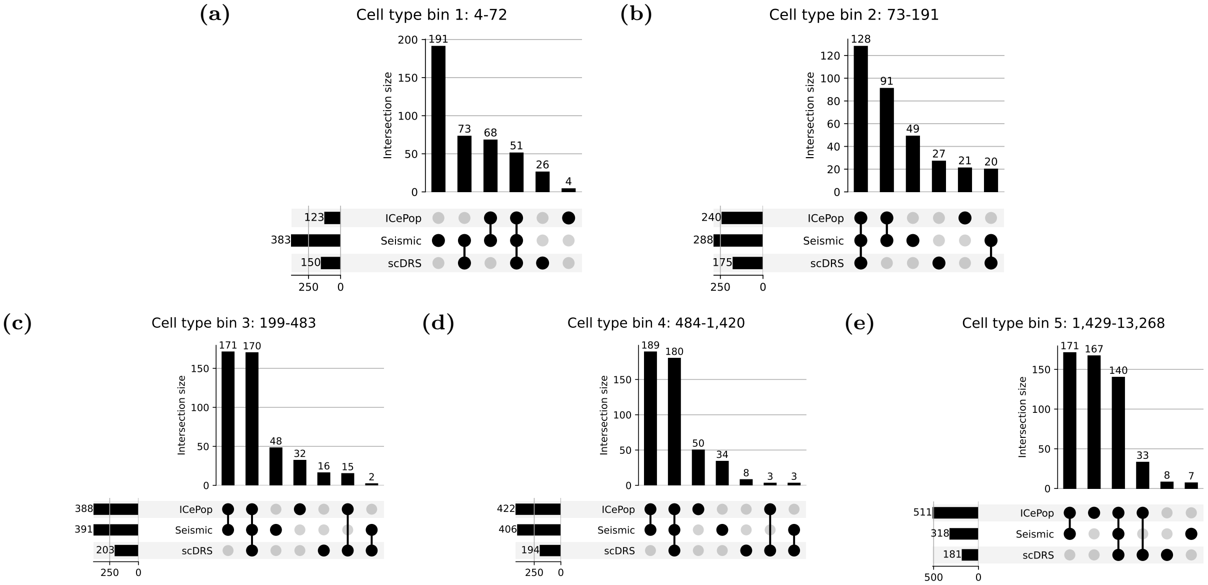


**Figure S11: Comparison of disease-cell-type associations among ICePop, seismic and scDRS across cell type sizes.** UpSet plot summarizing shared and unique disease-cell-type associations identified from the Tabula Muris FACS dataset across cell type size groups for ICePop, seismic, and scDRS. Cell type sizes are stratified into five quantile bins. In panels a-e, the bar plots on the left indicate the total number of associations detected within each size bin for three methods, while the top intersection bars show the number of shared and size-specific associations across methods.

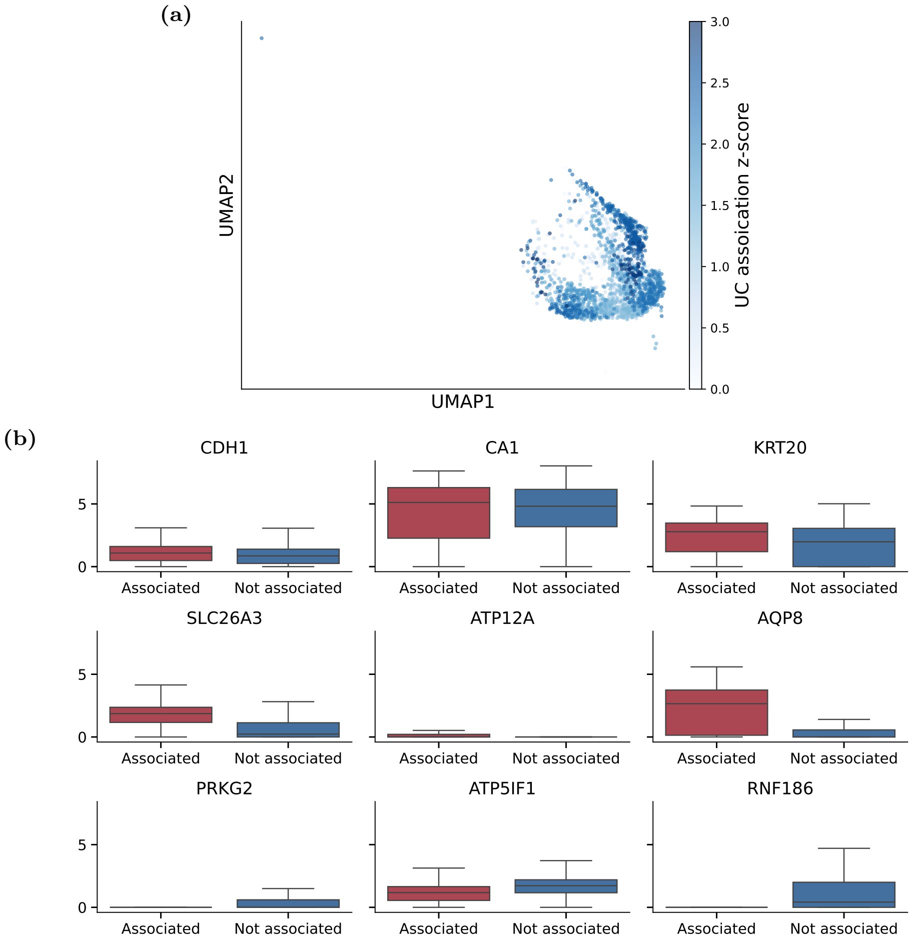


**Figure S12: Ulcerative colitis association in large-intestine enterocytes.** (a) UMAP showing ulcerative colitis (UC) association across large-intestine enterocytes. Metacell-level association scores were projected onto individual enterocytes. (b) Boxplots showing the expression of canonical large-intestine enterocyte marker genes (*CDH1*, *CA1*, *KRT20*), genes associated with differentiated enterocyte function (*SLC26A3*, *ATP12A*, *AQP8*), and genes involved in enterocyte homeostasis (*PRKG2*, *ATP5IF1*, *RNF186*). Red indicates UC-associated cells and blue indicates non-associated cells, defined by FDR $\leq 0.1$.

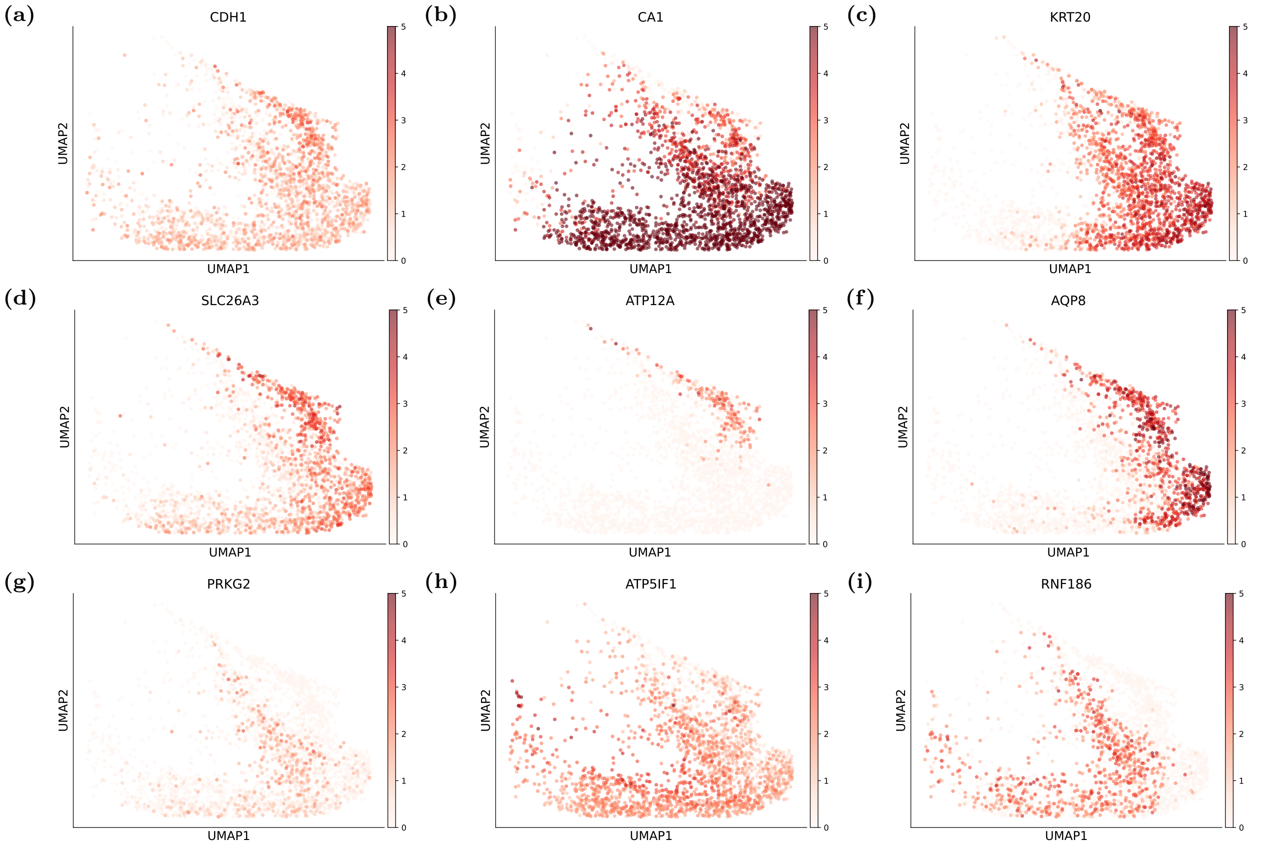


**Figure S13: Expression of marker and functional genes in large-intestine enterocytes.** UMAP showing the expression of canonical large-intestine enterocyte marker genes (panel a-c; *CDH1*, *CA1*, *KRT20*), genes associated with differentiated enterocyte function (panel d-f; *SLC26A3*, *ATP12A*, *AQP8*), and genes involved in enterocyte homeostasis (panel g-i; *PRKG2*, *ATP5IF1*, *RNF186*).

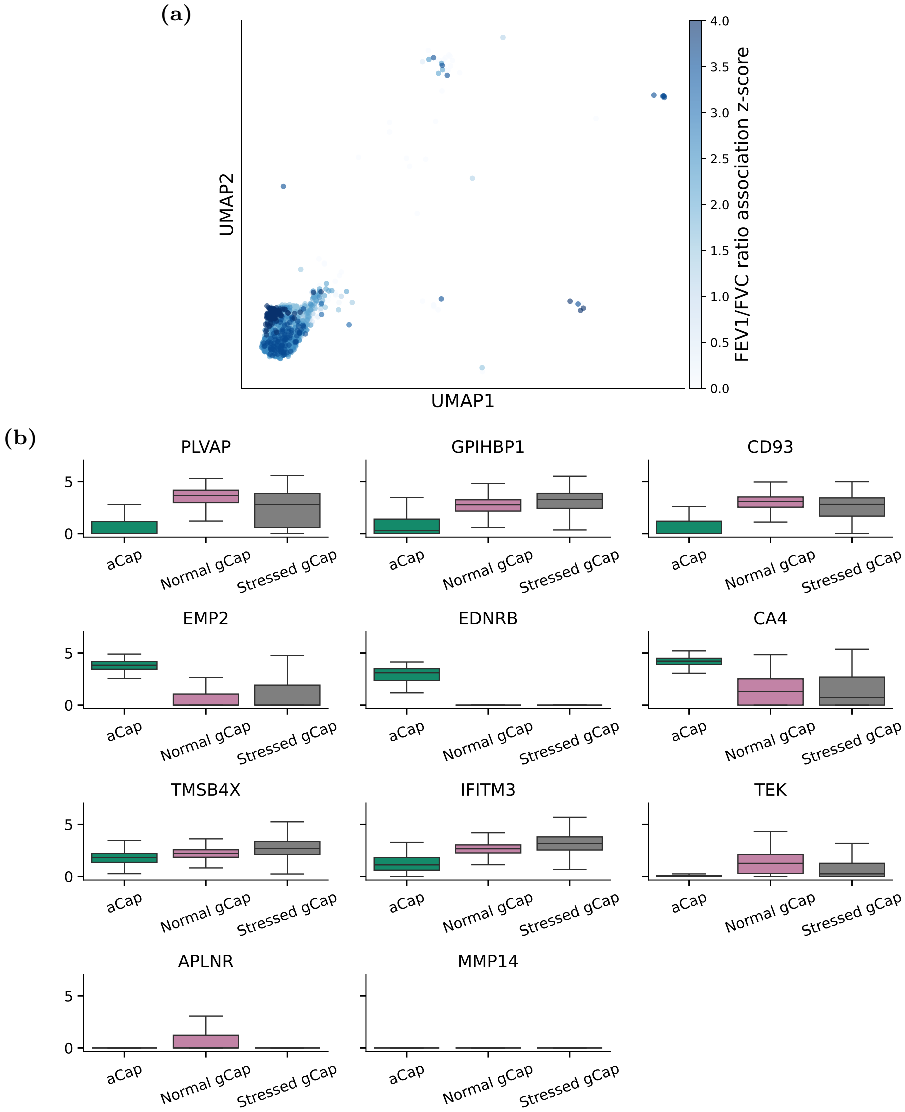


**Figure S14: FEV1/FVC ratio association in lung endothelial capillary cells.** (a) UMAP showing FEV1/FVC ratio association across lung endothelial capillary cells. Metacell-level association scores were projected onto individual cells. (b) Boxplots showing the expression of canonical general capillary (gCap) marker genes (*PLVAP*, *GPIHBP1*, *CD93*), canonical aerocyte (aCap) markers (*EMP2*, *EDNRB*, *CA4*), and genes associated with immune stress (*TMSB4X*, *IFITM3*). Genes reported to lose gCap identity under immune stress in gCap cells (*TEK*, *APLNR*, *MMP14*) are also shown.

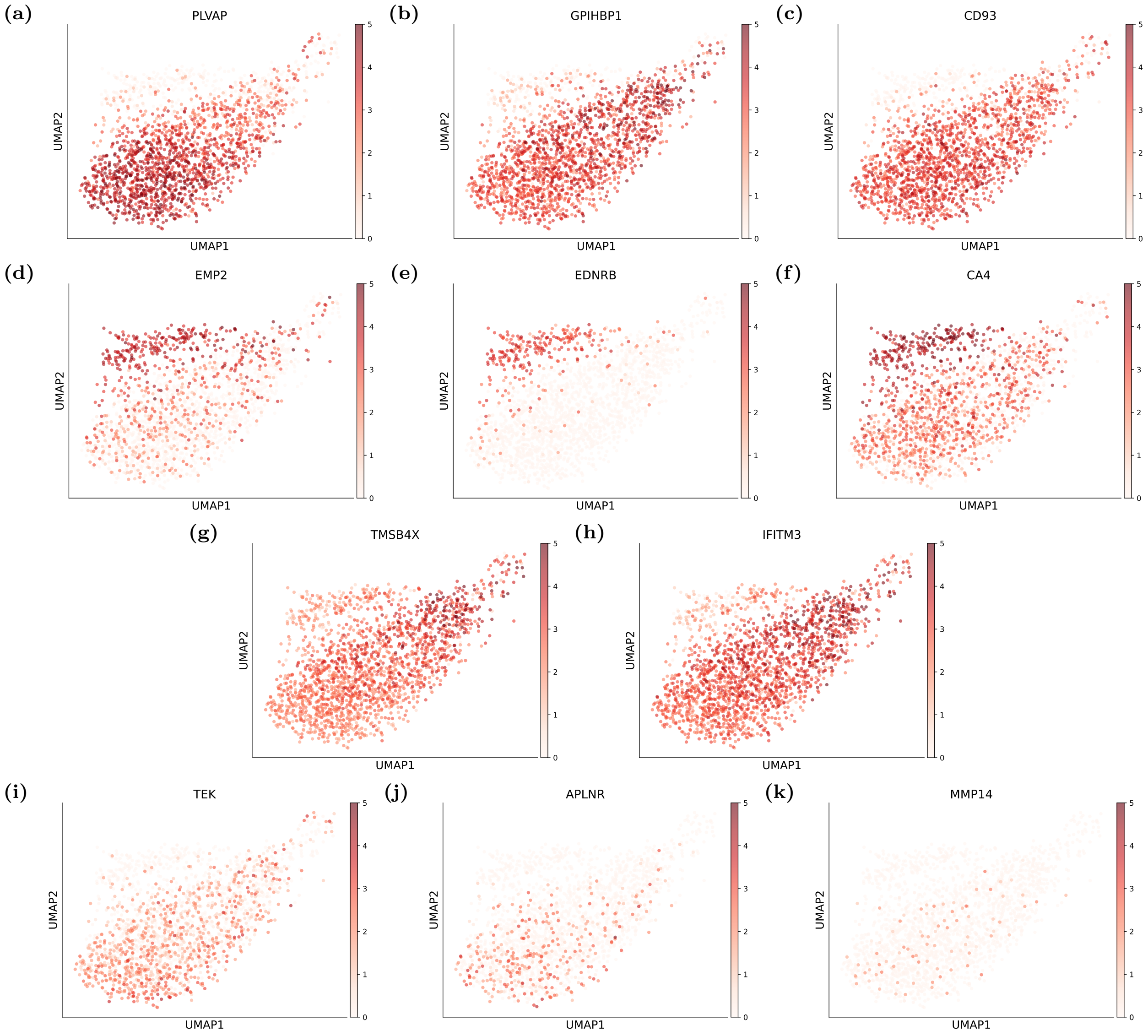


**Figure S15: Expression of markers in endothelial lung capillary cells.** UMAP showing the expression of canonical general capillary (gCap) marker genes (panel a-c; *PLVAP*, *GPIHBP1*, *CD93*), canonical aerocyte (aCap) markers (panel d-f; *EMP2*, *EDNRB*, *CA4*), and genes associated with immune stress (panel g-h; *TMSB4X*, *IFITM3*). Genes reported to lose gCap identity under immune stress in gCap cells (panel i-k; *TEK*, *APLNR*, *MMP14*) are also shown.

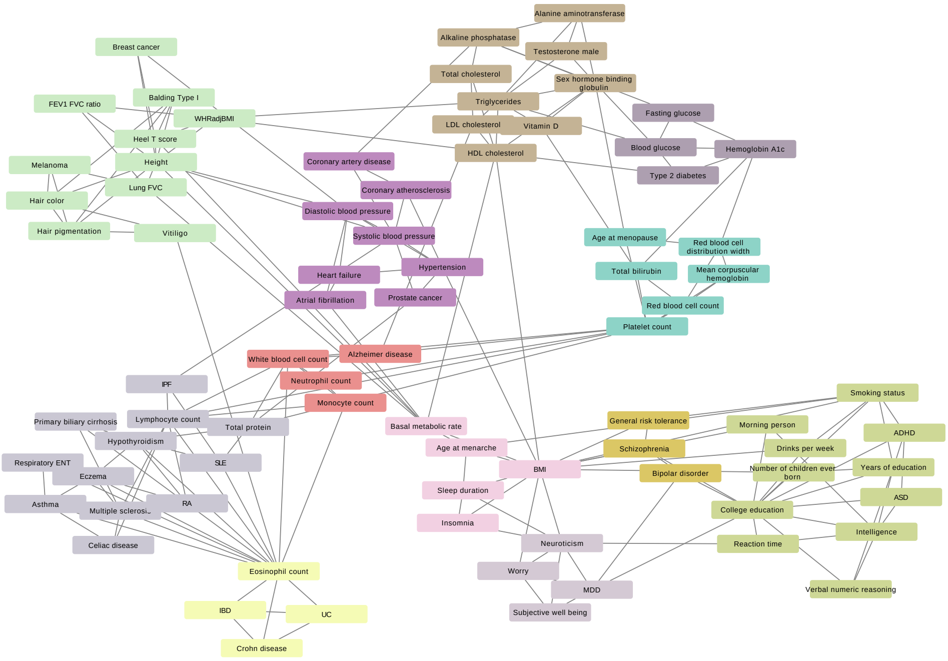


**Figure S16: Disease-disease similarity graph based on MAGMA z-scores, highlighting shared genetic risk.** The graph is constructed from similarities among quantile normalized MAGMA z-scores using a k-nearest-neighbor graph, where k=3. Louvain clustering is applied at a resolution of 2.5, with each color representing a distinct cluster of traits or diseases.

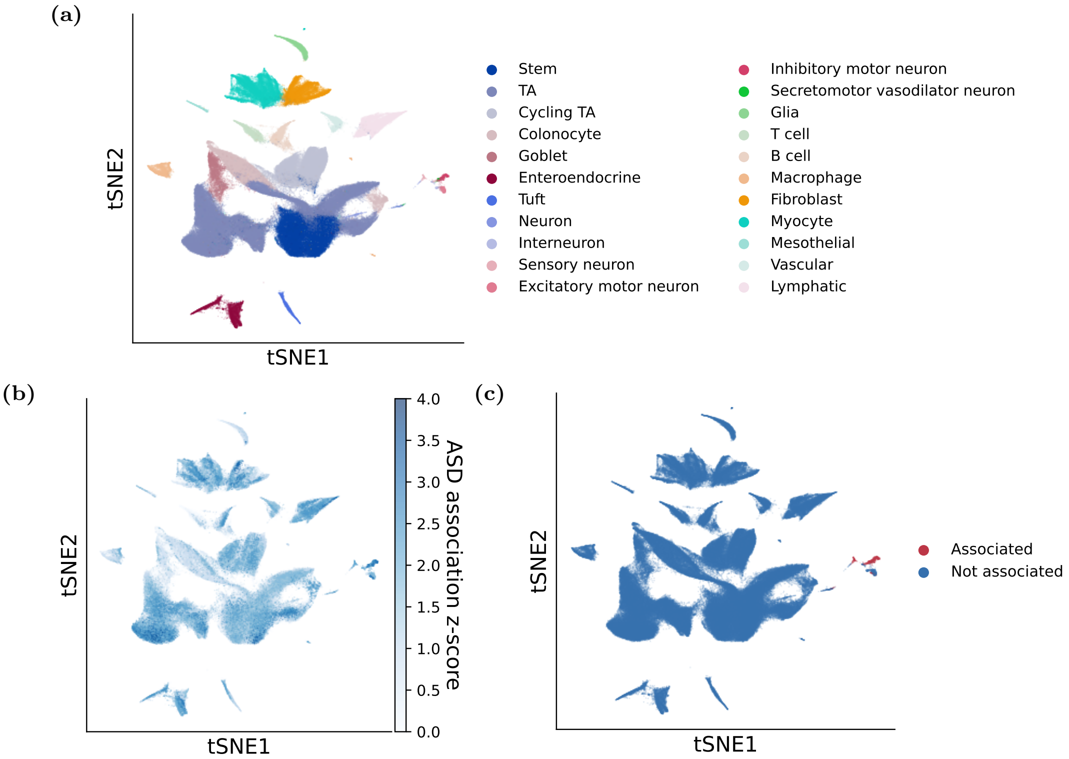


**Figure S17: ASD association in the mouse colon dataset.** tSNE of the mouse colon dataset showing (a) annotated cell types, (b) ICePop metacell-level ASD association scores projected onto individual cells, and (c) binarized association status (associated vs. non-associated), defined by FDR $\leq 0.1$.

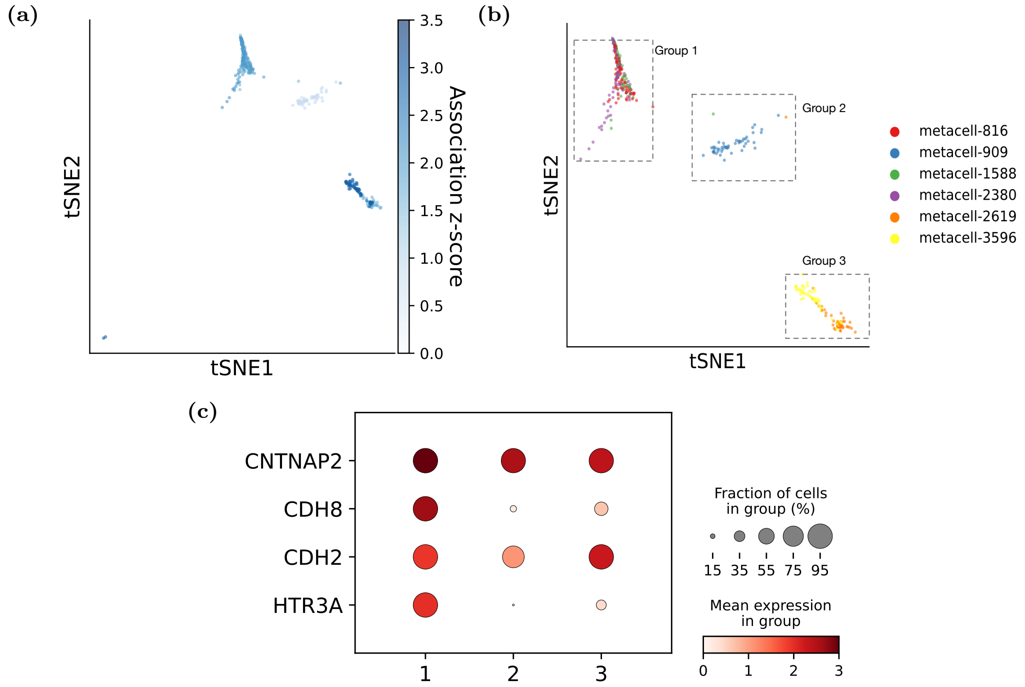


**Figure S18: Heterogeneity of ASD association in enteric sensory neuron subgroups.** tSNE of enteric sensory neurons showing metacell-level ASD association scores projected onto individual cells (a), and metacell distribution (b), with corresponding associated cell groups highlighted. (c) Dotplot showing the expression of genes known to be directly associated with dysfunction of enteric sensory neurons across three cell groups. Color indicates expression level, and circle size represents the proportion of cells expressing each gene.

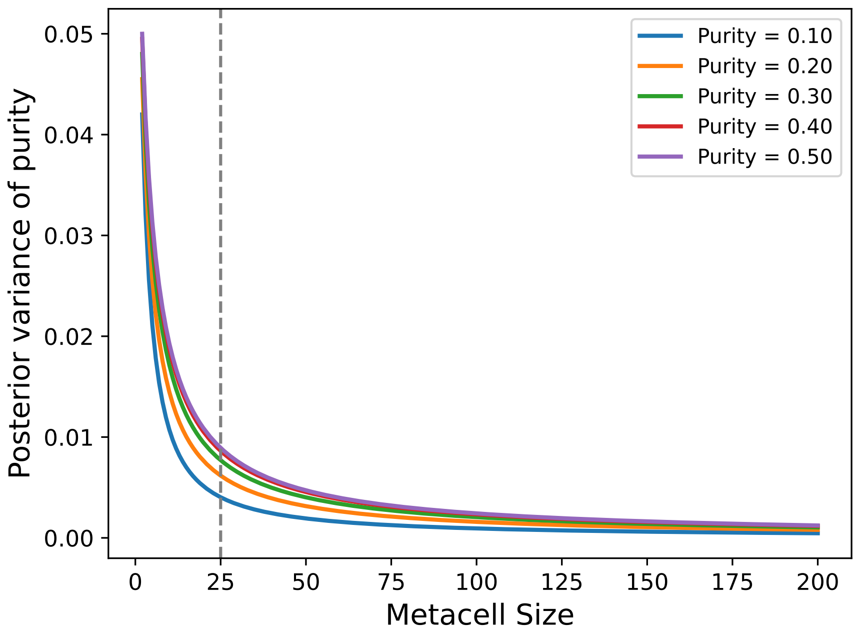


**Figure S19: Effect of metacell size on posterior uncertainty in purity.** The plot shows the posterior variance of cell type purity under a Beta-Binomial model. The x-axis represents metacell size, and the y-axis shows the posterior variance. Colors indicate different underlying purity levels. The dashed gray vertical line at metacell size 25 marks an elbow point where uncertainty decreases substantially.

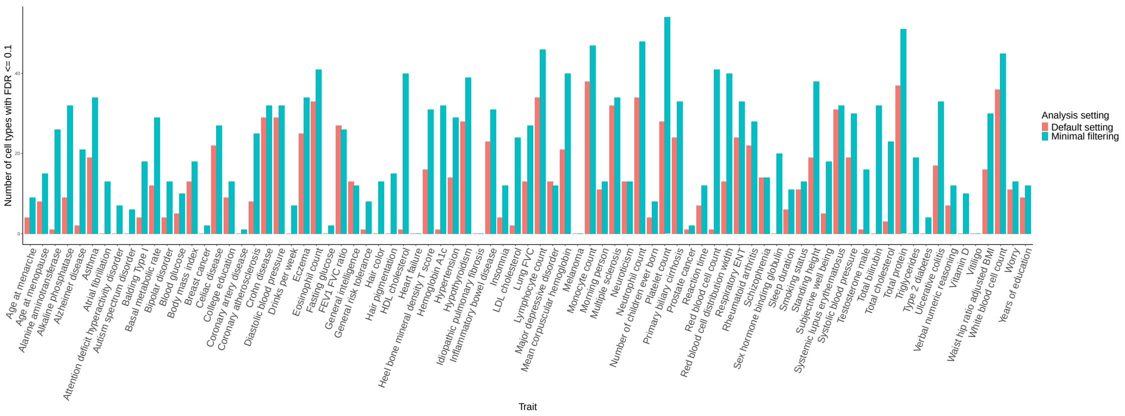


**Figure S20: Impact of default seismic filtering parameters on the number of significant cell types per trait.** Histograms show the distribution of the number of significantly associated cell types across traits under two filtering settings: the default average cell-type-level expression threshold ($min\_avg\_exp\_ct$ = 0.1, red bar) and minimal filtering with the threshold set to 0 (blue bar). A cell type is considered significant if FDR $\leq$ 0.1

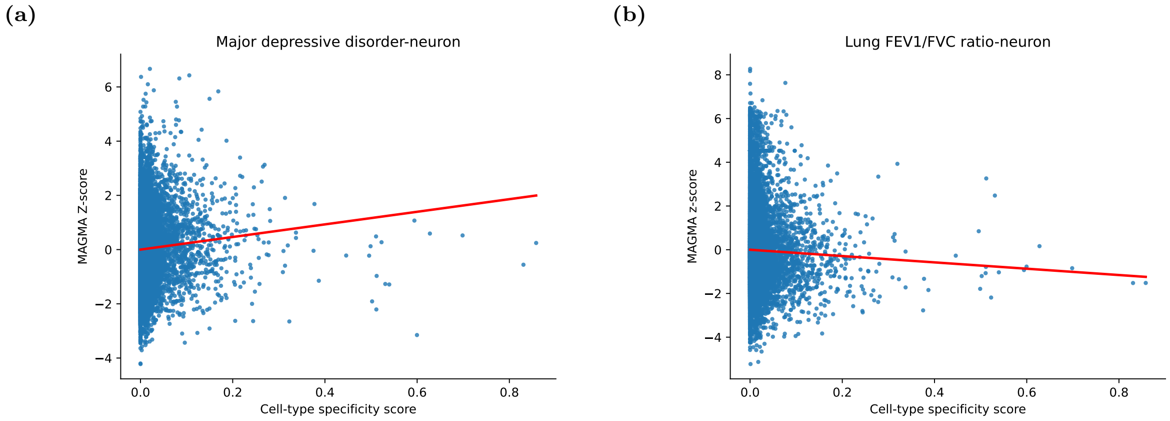


**Figure S21: Examples of positive and negative disease-cell type associations identified by seismic.** Scatter plots showing representative examples of a positive association between neurons and major depressive disorder (a) and a negative association between neurons and the FEV1/FVC ratio (b). The x-axis denotes the seismic specificity score, and the y-axis shows the corresponding MAGMA z-score. The red line represents the fitted regression line.

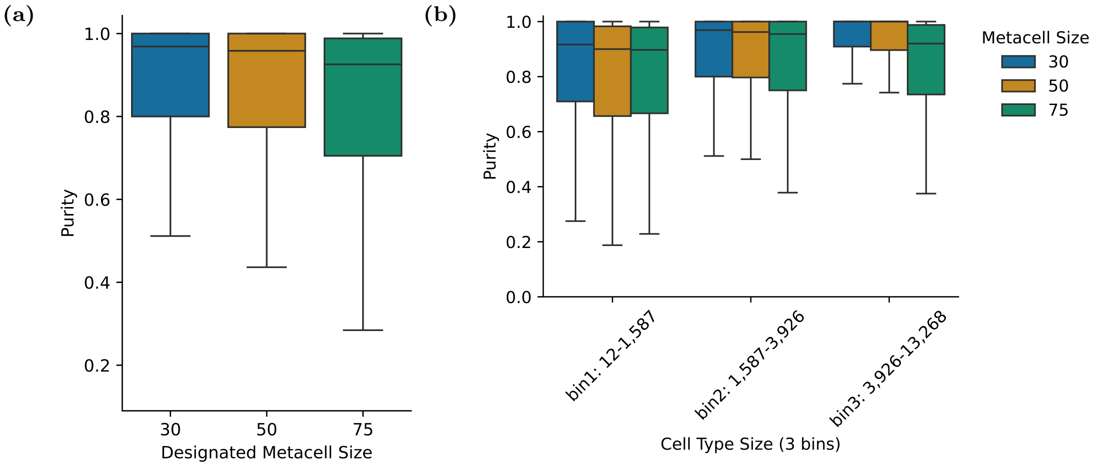


**Figure S22: Cell-type purity distribution for the Tabula Muris FACS dataset across different metacell sizes.** (a) Boxplots show the distribution of overall cell-type purity for metacell sizes of 30, 50, and 75. (b) Purity distribution are further stratified across three cell-type-size bins.

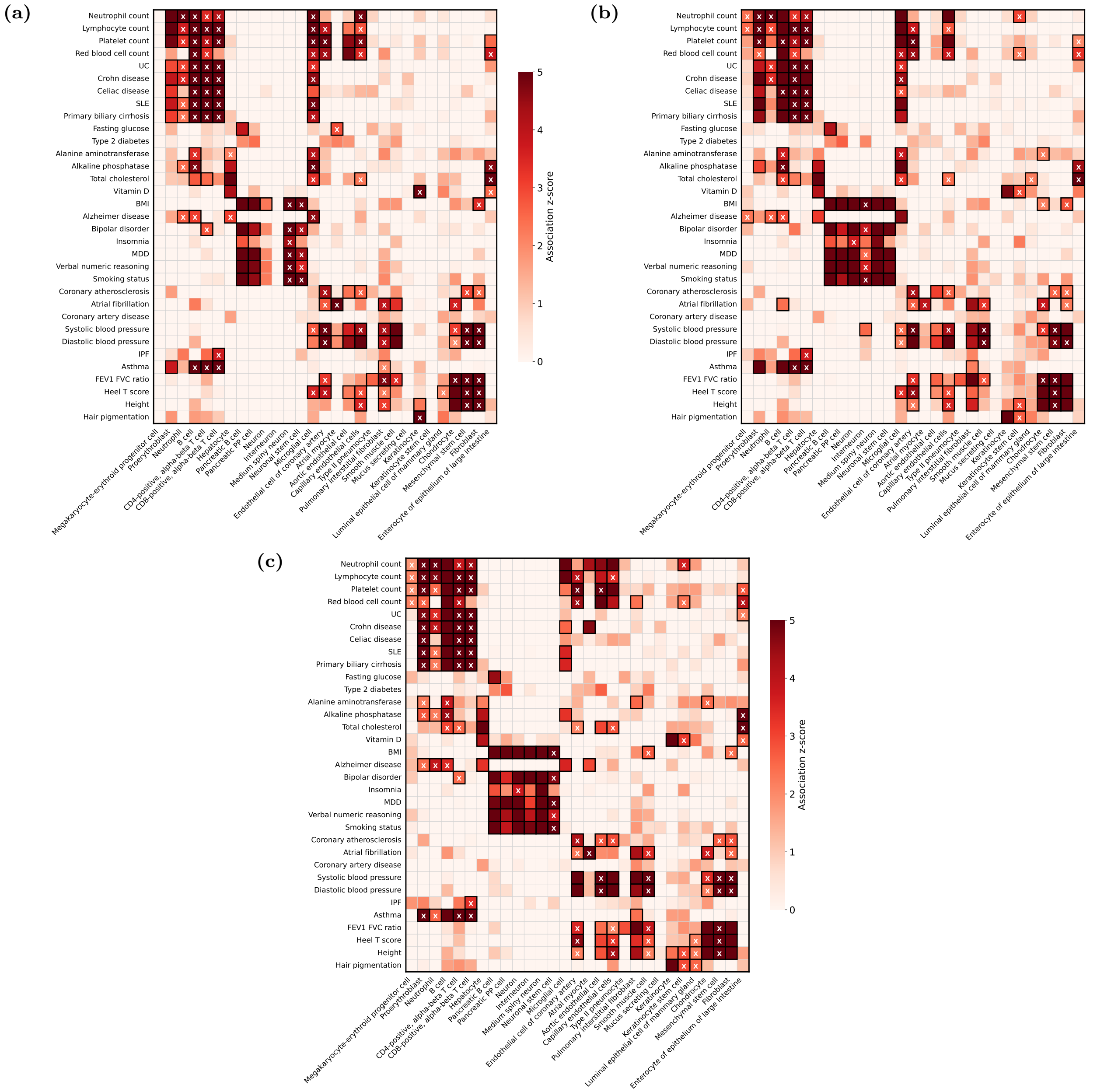


**Figure S23: Heatmap of ICePop disease-cell-type associations aggregated from three metacell sizes for 33 representative traits and 29 cell types from the Tabula Muris FACS dataset.** Rows correspond to diseases or traits, and columns correspond to cell types, grouped into seven disease categories and seven tissues or organs. Colors indicate cell-type association z-scores derived from ICePop, with negative values truncated to zero. Black rectangles denote significant associations, and cross symbols indicate heterogeneous associations, defined as significant signals (FDR $\leq$ 0.1) present in more than 20% but fewer than 80% of cells within a cell type. Trait abbreviations are ulcerative colitis (UC), systemic lupus erythematosus (SLE), body mass index (BMI), major depressive disorder (MDD), and idiopathic pulmonary fibrosis (IPF). Panels a-c show association results for metacell sizes of 30, 50, and 75, respectively. Association results for all 81 traits and 120 cell types across metacell sizes are provided in Supplementary Data 2.

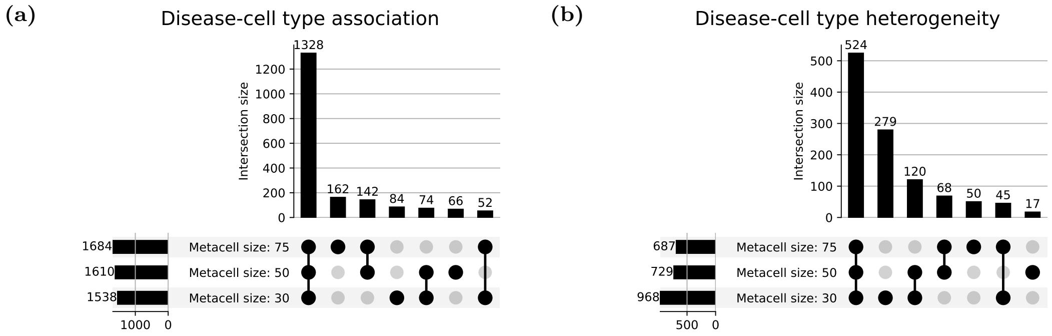


**Figure S24: Comparison of disease-cell-type associations across metacell sizes.** (a) UpSet plot summarizing shared and unique disease-cell-type associations identified from the Tabula Muris FACS dataset across metacell sizes. (b) UpSet plot summarizing shared and unique identification of heterogeneous associations among associations shared across all three metacell sizes. Heterogeneity is defined as a significant signal present in more than 20% but fewer than 80% of cells within a cell type. In both panels, the left bar plots indicate the total number of disease-cell type associations (or heterogeneous associations) detected for each metacell size, while the top intersection bars show the number of shared and size-specific disease-cell type associations (or heterogeneous associations).

#### Supplementary Tables

| Cell type | P | FDR | Method |
|---|---|---|---|
| Inhibitory motor neuron | 0.0053 | 0.0768 | ICePop |
| Secretomotor vasodilator neuron | 0.0105 | 0.0768 | ICePop |
| Sensory neuron | 0.0101 | 0.0768 | ICePop |
| Secretomotor vasodilator neuron | 0.0016 | 0.0351 | seismic |

Table 1: Significant ASD-cell type association in mouse colon among ICePop, seismic and scDRS (FDR: false discovery rate). Significant associations are detected by ICePop and seismic, but not by scDRS.

### Supplementary Notes

### 1 Related works

#### seismic

Seismic [1] identifies disease-associated cell types by first computing gene specificity scores. Gene specificity is defined by comparing the expression of a focal cell type to the remaining cell types. This specificity score is further combined with the expression coverage of each gene within the focal cell type and then normalized across cell types to derive the seismic score. Disease-cell-type associations are then quantified by regressing seismic scores against MAGMA gene z-scores. A positive association indicates that genes highly specific to a given cell type tend to have higher MAGMA z-scores, whereas a null or negative association indicates that cell-type-specific genes are either randomly distributed across MAGMA z-scores or enriched among genes with lower MAGMA z-scores.

Conceptually, seismic does not directly test for discrete "positive", "negative" or "no" enrichment. Rather, it evaluates whether the distribution of cell-type-specific genes is skewed toward higher, lower MAGMA z-scores or randomly distributed. We illustrate this with a clear positive example: major depressive disorder and neurons, where neuron-specific genes are enriched among genes with high MAGMA z-scores (**Supplementary Fig. S21a**). Conversely, a negative example is observed for the association between lung FEV1/FVC ratio and neurons, where neuron-specific genes are concentrated among genes with lower MAGMA z-scores (**Supplementary Fig. S21b**). Because this framework measures overlap between cell-type-specific genes and disease-associated genes through a regression-based enrichment model, it captures graded shifts in distribution. This makes Seismic sensitive to detecting cell-type populations with modest but consistent signals.

Seismic adopts the DFBETAS framework [2] to identify genes that contribute disproportionately to disease-cell-type associations. Specifically, DFBETAS quantifies the influence of each gene on the regression between seismic scores and MAGMA z-scores. Genes that drive the association upward, i.e., those with both high seismic specificity scores and high MAGMA z-scores—tend to have large positive DFBETAS values. An empirical threshold of $\frac{2}{\sqrt{n}}$, where $n$ is the number of genes included in the regression, is used to identify influential genes. However, genes with extremely high specificity scores but only moderate MAGMA z-scores may still exceed this threshold. A more stringent cutoff (e.g., $\frac{3}{\sqrt{n}}$) or restricting attention to the top-ranked influential genes may yield a more biologically meaningful set of candidates.

#### scDRS

scDRS [3] infers disease-cell-type associations by first computing disease scores at the individual cell level. By default, scDRS selects the top 1,000 genes ranked by MAGMA gene-level z-scores as the disease gene set. For each cell, it tests whether these disease-associated genes are overexpressed relative to matched control gene sets. The control gene sets are constructed via Monte Carlo sampling to match the disease gene set in size, mean expression and expression variance. For each cell, scDRS normalizes the set wise expression score across sets and cells, then computes a disease score and derives a cell-level p-value by comparing the disease set expression to the distribution of sampled control sets. To obtain cell-type-level associations, scDRS aggregates cell-level scores within each cell type. Specifically, it uses the 95th percentile (top 5%) of normalized disease scores within the cell type as the group-level statistic, and evaluates its significance using a Monte Carlo test based on the corresponding control scores. Although the choice of 5% may appear somewhat arbitrary, the authors show that results are robust to moderate changes in this threshold.

Because scDRS operates at single-cell resolution, it also enables testing for within-cell-type heterogeneity. To assess whether disease association signals cluster within subsets of cells, scDRS applies Geary's C, a spatial autocorrelation statistic, to determine whether high disease scores are non-randomly localized within the cell-cell neighborhood graph. This allows identification of heterogeneous subpopulations driving the association signal.

Notably, scDRS assumes that disease-associated genes, tend to show overall higher expression relative to matched control gene sets. This assumption can make the method conservative. In particular, some biologically important disease-related genes may not exhibit high expression but instead show highly specific expression in restricted cell populations. Because scDRS emphasizes relative overexpression compared to broadly matched controls, such specifically expressed but low-abundance genes may contribute less to the aggregated disease score. Together, these factors may reduce sensitivity and lead to conservative detection of disease-cell-type associations in certain contexts.

scDRS prioritizes disease-relevant genes by computing the correlation between the expression of each gene and the scDRS disease score for a given trait. Genes that show strong positive correlation with the disease score are considered co-expressed with GWAS-implicated genes and are therefore prioritized as candidate disease-associated genes. Notably, this correlation is computed across all cells in the dataset (e.g., the Tabula Muris FACS dataset used in the scDRS paper). As a result, the method tends to prioritize genes whose expression is broadly associated with the disease score across multiple cell types. In contrast, genes that are strongly associated with the disease score in only a small subset of cell types may be diluted in the global correlation analysis and result in lower correlation.

### 2 Adjustment of seismic's filtering parameters

Seismic filtering criteria can drastically affect the number of resulting predicted associations. We found that the default filtering parameters were too strict and left us with too few genes passing specificity filtering, which in turn led to very few downstream associations.

In particular, the *min_avg_exp_ct*, which is average expression of genes across cell types, set at threshold of 0.1 was overly restrictive in our testings. Although this parameter is intended to prevent specificity scores from being affected by very low average expression within a cell type, in practice it removed a large number of genes. We tested this on the Tabula Muris FACS dataset and found that the filtering criteria reduced the number of genes from 22,124 to 6,163 after specificity score filtering.

While originally introduced to stabilize the specificity score, the *min_avg_exp_ct* threshold substantially reduced the size of the gene set. To retain sufficient power for downstream analyses, we decided to set *min_avg_exp_ct* to zero. This minimal filtering approach recovered substantially larger number of disease-associated cell types (**Supplementary Fig. S20**). Among these additional associations, we observed biologically meaningful signals, including discovery of immune cell types (e.g., promonocyte, myeloid dendritic cell, monocyte) associated with Rheumatoid Arthritis (**Supplementary Data 2**).

### 3 Choice of metacell size

To evaluate the impact of metacell size, we systematically examined smaller sizes (30 and 50) in both simulation and real datasets. In simulation analyses, metacell size had no detectable effect on null performance (**Supplementary Fig. S1, S2, and S3**) and did not influence statistical power when varying the fraction of causal genes, noise levels, or signal scaling factors (**Supplementary Fig. S5a-c and S6a-c**). However, under heterogeneous sampling scenarios, where cell type heterogeneity level varied, smaller metacell sizes yielded slightly higher power (**Supplementary Fig. S5d and S6d**). Nonetheless, the absolute differences in power were modest.

TM analyses revealed a clearer trade-off between detection power and heterogeneity resolution across metacell sizes. We observed that the three tested metacell sizes (30, 50 and 75) produced generally comparable distributions of cell type purity, with 75 have slighly lowered purity (**Supplementary Fig. S22**). Among significant disease-cell type associations across all metacell size, the smallest metacell size (n=30) yielded the greatest number of unique heterogeneity detections (n=186) (**Supplementary Fig. S24b**), consistent with cases such as microglia in Alzheimer's disease, where heterogeneity was resolved only at this finer resolution, while larger metacell sizes (50 and 75) captured these associations as homogeneous (**Supplementary Fig. S23a**). However, the largest metacell size produced the greatest number of unique overall detections (n=162), substantially exceeding those observed at intermediate sizes (n=66 for size 50 and n=84 for size 30) (**Supplementary Fig. S24a**), as illustrated by broader associations between interneurons and neurological diseases that were detected only with larger metacells (**Supplementary Fig. S23b and c**). These results suggested that metacell size can be tuned according to analytical priorities: larger metacells improve broad detection power, while smaller metacells enhance the resolution of intra-cell type heterogeneity among strongly associated disease-cell type pairs. In this work, we focused primarily on balancing these objectives.

In most of our analysis, we selected an expected metacell size of 75 cells, a heuristic within the empirically stable range reported by Persad et al. [4], where SEACells demonstrated robust performance in representing distinct cell types, minimizing non-trivial multiple metacell assignments, and preserving rare cell types as high-purity metacells. Consistent with our results, this choice provides strong overall detection power and robustness, while smaller metacell sizes may be preferred when the primary goal is to resolve finer intra-cell type heterogeneity.

### 4 Examples of cases missed by ICePop but revealed in seismic

Despite advantages of ICePop, we observed instances in which other methods detected biologically meaningful associations that were not significant under ICePop. For example, respiratory basal cells are known to be affected in asthma, particularly due to abnormal basal cell differentiation [5]. Seismic identified a significant association for asthma in respiratory basal cells (p=0.024, FDR=0.088), whereas ICePop did not reach significance (p=0.087, FDR=0.275) (**Supplementary Data 3**). This discrepancy may reflect slightly reduced statistical power in metacell-based aggregation in certain contexts. Nevertheless, the aggregated z-scores in ICePop still suggested a related signal.

### 5 Limitation of perturbing gene expression in null and causal simulation

Both scDRS [3] and seismic [1] evaluated the performance using simulations under null and causal scenarios. In these frameworks, target cells are randomly selected, signals are removed by permuting gene indices, and causal effects are introduced by perturbing gene expression by effect size. This simulation strategy has several limitations. First, randomly modifying gene expression directly alters the underlying expression landscape in a brute-force manner, producing patterns that are unlikely to reflect realistic biological variation. Second, defining causal cell populations by randomly sampling cells across the dataset results in groups whose expression profiles are inherently heterogeneous and therefore do not resemble genuine cell types. Third, partitioning cells into only "causal" and "non-causal" groups is biologically unrealistic. In such settings, neither group follows a real cell-type expression distribution. Moreover, existing methods involve normalization across cell types. For example, seismic normalizes the product of cell-type specificity probability and expression coverage of a gene to sum to one across cell types. When normalization is performed across only two groups (causal vs. non-causal), the normalized scores and their differences are artificially inflated relative to realistic scenarios. This leads to overly strong signals and creates an unfair advantage over methods that construct signals at the individual cell or metacell level, then aggregate across cells or metacells to get cell type-level signals. To address this limitation, we instead perturb disease association signals at the level of MAGMA gene-level scores. This strategy preserves the full cellular structure and maintains the intrinsic expression continuum within each cell type, resulting in simulations that better reflect realistic biological scenarios.

### References

[1] Lai, Q., Dannenfelser, R., Roussarie, J.-P. & Yao, V. Disentangling associations between complex traits and cell types with seismic. *bioRxiv* (2024).

[2] Belsley, D. A., Kuh, E. & Welsch, R. E. *Regression diagnostics: Identifying influential data and sources of collinearity* (John Wiley & Sons, 2005).

[3] Zhang, M. J. *et al.* Polygenic enrichment distinguishes disease associations of individual cells in single-cell rna-seq data. *Nature genetics* **54**, 1572–1580 (2022).

[4] Persad, S. *et al.* Seacells infers transcriptional and epigenomic cellular states from single-cell genomics data. *Nature biotechnology* **41**, 1746–1757 (2023).

[5] Raby, K., Michaeloudes, C., Tonkin, J., Chung, K. & Bhavsar, P. Mechanisms of airway epithelial injury and abnormal repair in asthma and copd. front immunol 14: 1201658 (2023).
